# Polygenic outcomes of sexually antagonistic selection

**DOI:** 10.1101/2023.03.02.530911

**Authors:** Pavitra Muralidhar, Graham Coop

## Abstract

Sexual antagonism occurs when males and females have different fitness optima for a phenotype, but are constrained from evolving to these optima because of their shared genome. We study the response of a polygenic phenotype to the onset of sexually antagonistic selection, modeling a phenotype initially under stabilizing selection around an optimum, followed by a sudden divergence of the male and female optima. We observe rapid phenotypic evolution to these new optima via small changes in allele frequencies genome-wide. We study the role of sex chromosomes in this divergence and find that, in the absence of dosage compensation, the X chromosome favors evolution toward the female optimum, inducing co-evolutionary male-biased responses on the autosomes. However, dosage compensation obscures the female-biased interests of the X, causing it to contribute equally to male and female phenotypic change. In both scenarios, we see little effect of dominance in the genetic variation utilized by the X chromosome vs. the autosomes. We go on to examine the dynamics of stabilizing selection once the male and female optima have been reached, exploring a subtle mechanism through which the X chromosome, via the Bulmer effect, can cause higher equilibrium phenotypic variance in males than females. Finally, we consider how sexual antagonistic selection might persist across longer time scales, demonstrating that random fluctuations in an adaptive landscape can generate prolonged intragenomic conflict. Overall, our results provide insight into the response of complex phenotypes to sexually antagonistic selection and the evolution of sexual dimorphism.

## 1 Introduction

Sexually antagonistic selection arises when males and females have different fitness optima for a shared phenotype, but are hindered in evolving to these different optima because they share the majority of their genome. This form of intragenomic conflict is predicted to be common in any species with separate sexes, as males and females within a species will often experience different selection pressures due to their distinct life histories and reproductive strategies (Rice and Chippindale 2001; Chapman et al. 2003; Van Doorn 2009; Bonduriansky and Chenoweth 2009; Mank 2017a; Ruzicka et al. 2020). The fitness costs which sexually antagonistic conflict imposes on a population may eventually be resolved through a variety of evolutionary mechanisms (Wright et al. 2018; Mank 2017b; Connallon and Clark 2010, 2011) which lead to sexual dimorphism—quantitative or qualitative differences between males and females within a species.

Sexually antagonistic selection is therefore likely a key driver of intraspecific differentiation between males and females. Recognizing this, there is a considerable theoretical literature on the evolutionary dynamics of sexually antagonistic selection and its resolution. This literature can be divided into two general schools based on their distinct modeling approaches and assumptions.

The first school takes a ‘genotype-forward’ approach, and generally examines a single locus, or small set of loci, at which alleles carry fitness benefits in one sex but costs in the other (Haldane 1962; Kidwell et al. 1977; Patten and Haig 2009; Fry 2010; Jordan and Charlesworth 2012). The invasion probabilities of such alleles and the possible post-invasion dynamics may then be studied in detail. This approach has provided a tremendous amount of intuition for the circumstances under which sexually antagonistic selection can promote the maintenance of genetic variation, and has offered clear predictions for how alleles at major-effect loci will respond to sexually antagonistic selection—predictions that have subsequently been validated by the discovery of many examples of alleles (or sets of linked alleles) across different species that are maintained by sexually antagonistic selection (Roberts et al. 2009; Barson et al. 2015; Rusuwa et al. 2022). These genotype-forward analyses rely on the standard population genetic modeling approach of arbitrarily assigning selection coefficients (here, sex-specific selection coefficients) to individual alleles, without specifying details of the phenotype under selection (Ruzicka and Connallon 2022).

In contrast, the second school could be described as taking a ‘phenotype-forward’ approach, stemming from the quantitative genetic, rather than population genetic, tradition (Lande 1980; Turelli and Barton 2004; Wyman et al. 2013; De Lisle 2021). In this approach, it is assumed that an infinite—or extremely large—number of loci underlie variation in the phenotype, and that the effects of the alleles that segregate at these loci are approximately identical and very small. Under these assumptions, the evolutionary response of a genetically correlated male and female phenotype to a change in the male and female fitness optima can be studied in detail. This approach has provided substantial insight into expected patterns of phenotypic change under sexual antagonism, but is more limited in the predictions it can make about the allelic dynamics underlying that phenotypic change.

While both approaches have provided considerable insight into the dynamics of sexually antagonistic selection, recent genomic evidence suggests that the genetic architecture of most phenotypes is in fact polygenic, involving many loci of varying effect scattered throughout the genome. Motivated by these findings, our aim here is to study the response of a polygenic phenotype to sexually antagonistic selection, drawing upon both the ‘genotype-forward’ and ‘phenotype-forward’ approaches. In our model, the selective costs and benefits of individual alleles arise organically from a model of selection on the phenotype. This combination of approaches is facilitated by the recent development of whole-genome evolutionary simulation software (Haller and Messer 2019), enabling us to simultaneously study the phenotypic and genotypic response to sexually antagonistic selection.

Modeling sexually antagonistic selection on a polygenic phenotype may prove especially useful for the further development of methods to detect sexual antagonism in natural populations. Many tests have been developed to detect sexual antagonism. Some limit selection to one sex experimentally and study the fitness consequences for the other sex (Rice 1996); some measure the cross-sex correlation of fitness using related individuals or experimental sex reversal (Poissant et al. 2010; Chippindale et al. 2001); others have identified major-effect alleles maintained under balancing selection owing to opposing selective effects in the two sexes (Barson et al. 2015; Rusuwa et al. 2022). Recently, there has been an increased focus on developing tests to detect sexually antagonistic selection using population genomic data (Mank 2017a; Ruzicka et al. 2020). These population genomic tests have typically been based on predictions for allelic dynamics derived from ‘genotype-forward’ models (Ruzicka et al. 2020). However, the application of these methods to large scale population genomic data sets, particularly in humans, has yielded ambiguous results, with limited evidence for ongoing sexually antagonistic selection (Cheng and Kirkpatrick 2016; Kasimatis et al. 2019, 2021; Cheng and Kirkpatrick 2020; Bissegger et al. 2020). While this lack of signal could indicate a genuine lack of ongoing sexual antagonism, it could also be that the predictions of ‘genotype-forward’ single-locus models do not adequately describe the response to sexually antagonistic selection on a phenotype with a more complex genetic basis. Studying the genetic response of polygenic phenotypes to sexually antagonistic selection may thus provide guidance for how this common intragenomic conflict might be detected in population genomic data.

In the model we study, we assume that the phenotype in question is under stabilizing selection. There is abundant evidence that stabilizing selection is a common form of selection on phenotypes (Sanjak et al. 2018; Simons et al. 2018; Sella and Barton 2019), including phenotypes that have been shown to be involved in sexually antagonistic conflict in a number of species (Mank 2017a; Stulp et al. 2012; Sanjak et al. 2018; Prasad et al. 2007; Abbott et al. 2010). We examine a scenario in which males and females initially experience stabilizing selection around a common fitness optimum for the phenotype, but then the male and female fitness optima suddenly diverge. This divergence induces opposing directional selection on the phenotype in the two sexes (Lande 1976, 1980). During this phase of directional selection, alleles with concordant phenotypic effects in the two sexes will be beneficial in one sex and costly in the other. We examine the genetic and phenotypic response to the shift in male and female optima, drawing from an extensive theoretical literature on the response of polygenic phenotypes under stabilizing selection to shifts in fitness optima (de Vladar and Barton 2014; Jain and Stephan 2017; Thornton 2019; Stephan and John 2020; Hayward and Sella 2022).

Because sexually antagonistic selection reflects selection for divergent fitness optima in males and females, of especial interest are the consequences of this conflict for genomic regions that do not spend equal time in each sex. In particular, sex chromosomes (we focus on the X/Z chromosome) often comprise a significant portion of an organism’s genome, and therefore may have a substantial influence on the evolution of polygenic phenotypes. Sex chromosomes are predicted to have distinct evolutionary ‘interests’ relative to the autosomes, due to their asymmetric transmission patterns through males and females. Which sex is ‘favored’ by these distinct evolutionary interests has been a topic of considerable debate (Rice 1984; Haig 2006; Fry 2010; Frank and Patten 2020). However, this literature has tended to adopt the genotype-forward approach described above, focusing on individual autosomal and sex-linked loci of major effect, in isolation of each other. A major focus of our work is therefore not only to study the response of sex chromosomes to sexually antagonistic selection in an explicitly polygenic framework, but also to study how sex chromosomes co-evolve with the autosomes to resolve this intragenomic conflict between the sexes.

## 2 Methods

### Model overview

We study the response of a polygenic phenotype that initially experiences stabilizing selection around a common optimal value in both sexes to an instantaneous change in the fitness optima of males and females. For simplicity, we model only the genetic component of the phenotype, assuming no environmental effects on the phenotype (see Discussion). We first examine an additive genetic model, in which the phenotype of an individual is determined by summing the allelic effect sizes across all loci across both copies of the individual’s genome. In this case, an individual’s phenotype corresponds to their additive genetic value, or, in the context of the GWAS literature, their polygenic score. We allow mutations to occur at the loci underlying the phenotype, which generate new alleles that have an equal probability of increasing or decreasing the phenotypic value. We assume that the male and female effect sizes of each new allele are correlated, such that the effect size of a new mutation will often, though not always, be concordant in the two sexes (Fig. 1). This corresponds to a phenotype with a shared genetic basis between the sexes but also some level of sexual differentiation—a common situation for a variety of morphological and developmental phenotypes across taxa (Poissant et al. 2010; Kassam and McRae 2016).

**Figure 1:**
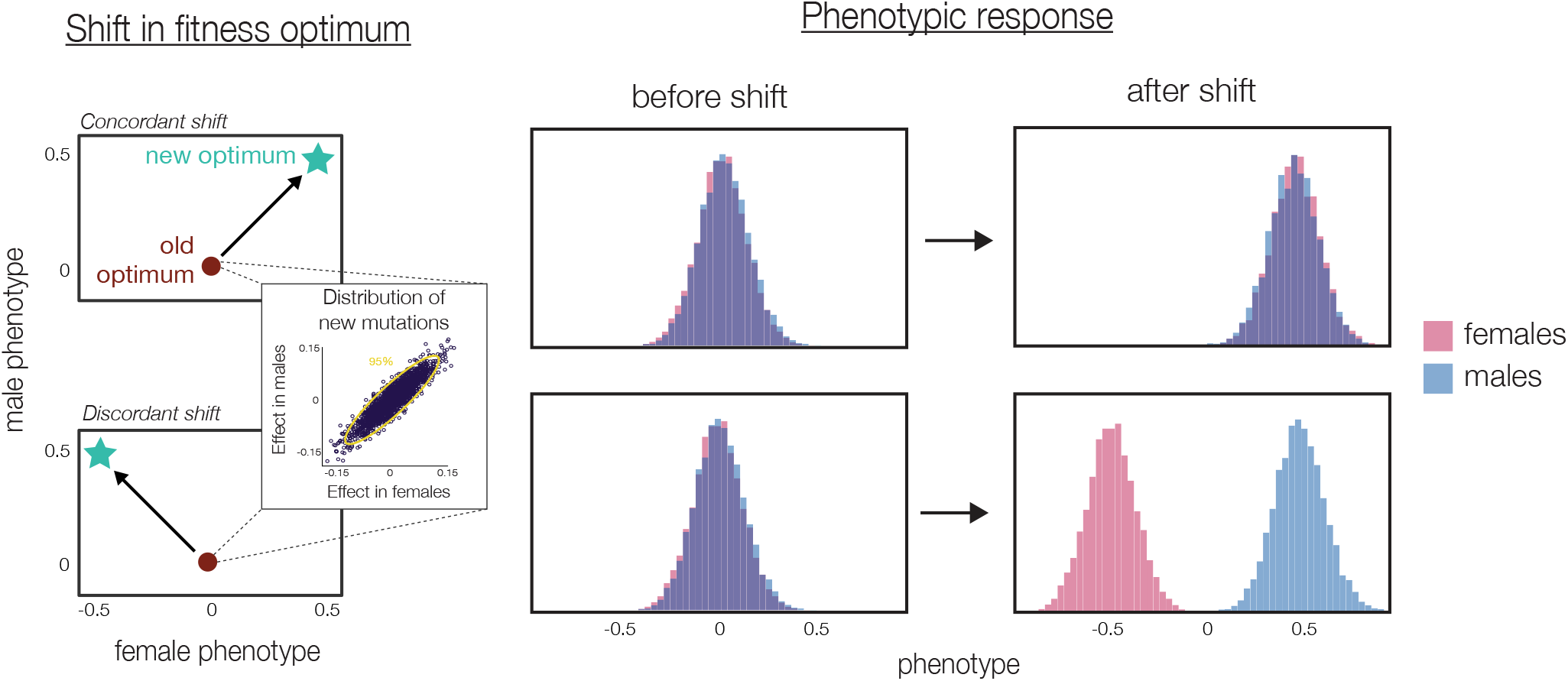
Divergence of male and female optima generates sexual antagonism. Illustrated are a concordant shift of the male-female optimum (top), in the same direction as the underlying male-female correlation of mutational effects on the phenotype (inset), and a discordant shift of the male-female optimum (bottom), orthogonal to the direction of the mutational correlation (inset). In both scenarios, the mean phenotype evolves toward the new male-female optimum. However, in the case of the discordant shift (bottom), movement to the new optimum involves opposing directional selection on the male and female phenotypes—that is, sexual antagonism. At the end of this process, once the population has reached the new male-female optimum, the male and female phenotypic distributions have diverged and sexual dimorphism in the phenotype has been established.

We simulate an instantaneous shift in fitness optimum in both males and females, so that the male optimum moves to an increased value, while the female optimum moves to a decreased value. This induces sexually antagonistic conflict, requiring males and females to evolve in opposite directions, against the underlying genetic correlation of the phenotype. As a contrast, we also consider the scenario where the male and female fitness optima suddenly shift in the same direction, so that the resulting phenotypic changes are aligned with the underlying genetic correlation.

### Simulation setup

The population is composed of 10,000 diploid individuals (5,000 male and 5,000 female throughout) and evolves according to a Wright-Fisher process with random mating. We implement stabilizing selection as a Gaussian fitness function with standard deviation *ω* = 1. Initially, the male and female fitness optima are identical: *O*_♂_ = *O*_♀_ = 0. After 100,000 generations, these optima shift such that *O*_♂_ = 0.5 and *O*_♀_ = −0.5. These correspond to a shift of approximately 2.5*VP*_0_ where *V*_*P*0_ is the phenotypic variance at the time of the shift. We then observe the population for 100,000 additional generations. The strength of stabilizing selection remains constant throughout.

The phenotype is controlled by 1000 loci (*L* = 1000). Unless otherwise stated, these loci freely recombine with each other, and there is no intralocus recombination. We implement a per-locus mutation rate of *u* = 10^-5^, so that the mutation rate per gamete per generation is *U* = *Lu* = 0.01. Unless otherwise stated, we assume that the effects of these mutations on the phenotype in males and females, *e_m_* and *e_f_*, are drawn from a bivariate normal distribution with variance 0.01 and a inter-sex correlation of 0.9. We also consider a scenario with reduced effect sizes, drawn from a bivariate normal distribution with variance 0.001. *e_m_* and ef describe the homozygous effect of an allele. The effect of an individual allele is therefore 0.5*e_m_* in males and 0.5*e_f_* in females in simulations in which additive effects are assumed, or *he_m_* and *he_f_* in simulations in which the dominance coefficient (*h*) varies. In simulations in which dominance varies, dominance coefficients were drawn from a uniform distribution between 0 and 1. We assume that the dominance coefficient of an allele is the same in males and females. At generation 0, we seed the population with 100 mutations, with effects drawn from the distribution described above and with frequencies chosen from a uniform distribution between 0 and 1.

After 100,000 (10*N*) generations, the fitness optima for males and females instantaneously diverge. We record the frequency and effect sizes of alleles segregating in the population at the time of the shift, and track the frequencies of these alleles across subsequent generations at different time points. Unless otherwise stated, we report the frequency of mutations that have appeared during the course of the simulation (i.e. derived alleles). We also record the average male and female phenotypic values at multiple time points before and after the shift in optimum.

All simulations were run across 10 replicates in SLiM 3.3 (Haller and Messer 2019). All analyses were conducted in R, and regression lines were calculated using the ‘lm’ function in the ‘stat’ package (R Core Team 2022).

### Single inheritance mode simulations

We initially study, in isolation, an entirely autosomal genome and an entirely X-linked genome, to clarify the evolutionary interests of these genomic regions. We have chosen to simulate an X chromosome rather than a Z chromosome, but all of our results equally apply to the Z chromosome (albeit with the sexes reversed).

In these simulations, we assume that our 1000 freely-recombining loci are all inherited according to the pattern of an autosome or an X chromosome. That is, in the case of an X, females will be diploid and males will be haploid, and males inherit all alleles from their mothers. We do not consider a sex-specific (Y) chromosome (see Discussion).

In our X chromosome simulations, we consider both the scenario where the X does not experience dosage compensation and the scenario where it does. Without dosage compensation, the effect on the male phenotype of an allele on the X is 0.5*e_m_* (regardless of the dominance of the allele in females); under dosage compensation, its effect on the male phenotype is *e_m_* (a proxy for the up-regulation of the entire X chromosome—but see Discussion).

### Full genome simulations

In these simulations, we consider a genome containing both sex chromosomes and autosomes. We simulate 1000 loci along a *Drosophila melanogaster*-like genome, using the linkage map produced by Comeron et al. (2012). Unless otherwise stated, we allow for autosomal recombination in males (unlike in *Drosophila*), which we assume to follow the autosomal map of female. We do not simulate a Y chromosome (but see Discussion). We assume that allelic effects at diploid loci are additive.

In the simulations, we separately tracked the additive genetic value for the phenotype derived from the X chromosome and autosomes, by summing up either the male or female effect sizes of all X-linked and all autosomal alleles carried by each individual.

### *F_ST_* calculations

Sexually antagonistic viability selection is expected to generate allele frequency differences between males and females within a generation. In order to calculate *F_ST_* between adult males and females, we had to slightly modify our basic simulations to explicitly incorporate viability selection (Wright-Fisher models in SLiM simulate selection as a decrease in the probability of mating, combining viability and reproductive selection). We calculated each individual’s phenotype at the start of each generation and calculated their expected fitness under our stabilizing selection scheme (which ranged between 0 and 1). We then drew a random number between 0 and 1, and if this number was greater than the individual’s fitness value, we excluded the individual from the sample over which we calculated *F_ST_*. We then calculated the average and maximum male-female *F_ST_*, using the formula set out in Cheng and Kirkpatrick (2016), across all segregating loci in this sample in the generation immediately following the shift in optima when sexually antagonistic selection is expected to be strongest. We repeat this process for 5 samples per replicate across 10 replication simulations.

### Moving optimum

We simulated a dynamic adaptive landscape in which the male and female fitness optima are continually changing. We assumed that the male-female optimum shifts every 500 generations by an absolute (Euclidean) distance of λ = 0.5. The direction of this shift is random, with its angle *A* drawn from a uniform distribution between 0 and 360 degrees. The new male and female optima are then 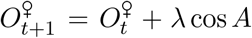 and 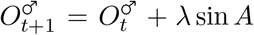. We tracked the angle at which the mean male-female phenotype moved every generation for 50,000 generations following the first optimum shift. In these simulations, we assumed a *Drosophila melanogaster*-like genome as described above.

## 3 Results

### 3.1 The sex chromosomes and autosomes, in isolation of each other, take different phenotypic paths to a new fitness optimum

We first study the phenotypic and genetic response of an entirely autosomal and an entirely X-linked genome to the divergence of the male and female fitness optima (Fig. 1). This shift induces opposing directional selection in males and females, driving the initial evolution of sexual dimorphism in the phenotype. Note that, while it is primarily to generate intuition that we initially study these special cases, they do in fact apply to haplodiploid species (for the X) and species with environmental sex determination (for the autosomes).

#### The phenotypic response

First, consider the case of an entirely autosomally-encoded polygenic phenotype. Because all individuals inherit a set of autosomes from their mother and their father, autosomes spend an equal amount of time in males and females, and will therefore respond equally to selection in males and females. This means that, assuming selection is acting equally strongly in males and females, the autosomally-encoded phenotype will take a straight path to the new optimum in male-female phenotype space (Fig. 2).

**Figure 2:**
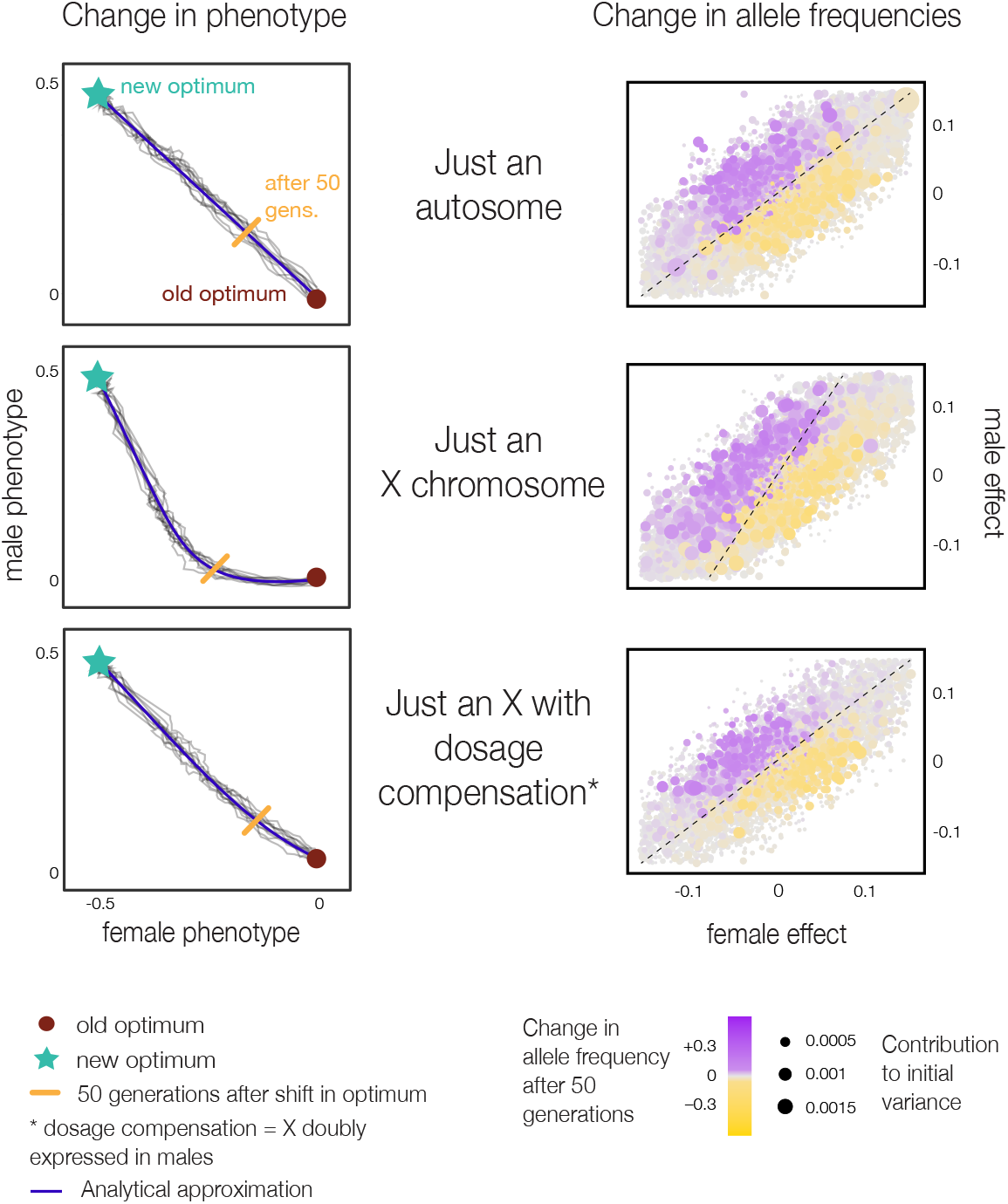
Autosomal and X-linked polygenic phenotypes take distinct paths to a new sexually antagonistic optimum. The left panels display the movement in male-female phenotype space of a polygenic phenotype encoded entirely on an autosome (top), an X chromosome without dosage compensation (middle), and a dosage-compensated X (bottom), following a discordant shift in the male-female fitness optimum. Black lines indicate the trajectories observed across 500 generations in each of 10 replicate simulations, while the solid indigo line shows the path predicted by the multivariate breeder’s equation modified for each case (Appendix). The gold bar indicates the mean phenotype 50 generations after the shift in optimum. Right panels show, for all alleles that were segregating at the time of the optimum shift, their changes in frequency across the subsequent 50 generations (color of bubbles) as a function of their effects in females (x-axis) and males (y-axis) and a proxy for their contribution to phenotypic variance at the time of the optimum shift (size of bubbles): 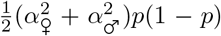 where *p* is the frequency of the allele. The dotted lines are the boundaries between alleles that are expected to be selectively favored vs. disfavored, such that alleles along the line should be selectively neutral. In the top and bottom panels, these dotted lines have slope 1, representing the equal weighting of male and female fitness for autosomes and for a dosage-compensated X chromosome; in the middle panel, the slope is 2, reflecting the female-biased interests of a non-dosage-compensated X chromosome.

In contrast, the X chromosome, due to its unique transmission pattern, spends twice as much time in females as in males. As a result, an X-encoded phenotype will respond twice as strongly to selection in females than in males, and will therefore initially evolve more rapidly toward the female optimum. As the mean female phenotype moves closer to its optimum, the strength of directional selection on the female phenotype will relax and directional selection on the male phenotype will begin to be relatively more prominent, resulting in eventual movement upward in male-female phenotype space toward the male optimum. This initially female-biased phenotypic response results in a characteristic curved path across the male-female phenotypic landscape toward the new optimum (Fig. 2). We see a somewhat similar trajectory for an X-encoded phenotype even in the case of a concordant shift in the male-female optimum, although the curvature of the phenotypic path is more extreme in the case of a sexually antagonistic shift (Fig. S6).

The X chromosome responds more strongly to selection in females because it spends twice as much time in females. However, many species have diverse mechanisms to equalize expression of the X between males and females despite its different copy number in each sex. Dosage compensation can occur through a variety of mechanisms, and its degree can also vary across the X chromosome itself (Mank 2013; Gu and Walters 2017). Here, we have chosen to implement dosage compensation by doubling the effect size of alleles on the X chromosome in males (Kent Jr et al. 2005). This is analogous to the dosage compensation mechanism observed in species like *Drosophila melanogaster,* where the expression of the X chromosome in males is doubled (Conrad and Akhtar 2012; Lucchesi and Kuroda 2015). This model also provides a good fit to X chromosome genetic variation for polygenic phenotypes in humans (Sidorenko et al. 2019). This form of dosage compensation increases the phenotypic variance of males, increasing the rate at which the X moves towards the male optimum. This approximately cancels out the effect of the X spending twice as much time in females, so that a polygenic phenotype encoded on a dosage-compensated X will take a path to the optimum similar to one encoded autosomally (Fig. 2; see also Charlesworth et al. 1987).

The response of correlated phenotypes to directional selection can be described via the multivariate breeder’s equation (Lande 1980; Walsh and Blows 2009), which can therefore describe the response of male and female phenotypes to selection despite their shared genetic basis (Cheng and Houle 2020). The multivariate breeders equation can also be modified to describe the response to selection of a X-encoded phenotype in males and females (Lande 1980; Fernando and Grossman 1990; Kent Jr et al. 2005; Yang et al. 2011) (Appendix). While this approach relies on assumptions of constant additive genetic variance and covariance across time, we find that these multivariate breeders equations, appropriately parameterized, provide an excellent approximation of the phenotypic response of a polygenic phenotype encoded entirely on autosomes or an X chromosome (Fig. 2; Appendix).

#### The genetic response

In our simulations, we can also characterize the allelic dynamics that underlie the divergent evolution of the male and female phenotype. Alleles may be divided into three categories based on their directional effects on the phenotype. First, those that decrease the phenotype in females and increase the phenotype in males. After the shift in optimum, these alleles are beneficial in both sexes, as they move both males and females closer to their respective fitness optima. There are also alleles which increase the phenotype in females and decrease it in males—these are deleterious in both sexes. The final category of alleles— those which increase or decrease the phenotype in both sexes—would traditionally be considered sexually antagonistic in genotype-forward models: they will be selected for in one sex and against in the other (Zhu et al. 2022).

Given this categorization, we can first examine the genetic response of an autosomally-encoded phenotype. The divergence of the male and female phenotype is caused by an increase in frequency of alleles in the first category described above, and a decrease in frequency of alleles in the second category (Fig. 2). We do not observe consistent changes in the frequency of alleles in the third category. These results are robust to decreasing the average effect size of individual alleles (Fig. S5).

In the case of the X, we observe different allelic dynamics. A polygenic phenotype encoded on a non-dosage compensated X chromosome responds more strongly to selection in females than in males, initially moving more quickly toward the female optimum. This is achieved by the increase in frequency of alleles that are aligned with the optimum shift in males and females, but also by those which decrease the phenotype in both sexes—that is, sexually antagonistic alleles that are beneficial in females and costly in males (Fig. 2). This pattern is consistent with previous analytical work, which has suggested that the X chromosome is likely to have female-biased ‘interests’ in the case of mainly additive alleles underlying a polygenic phenotype (Rice 1984; Fry 2010; Frank and Patten 2020). Another way to describe this pattern is that the marginal effect of an X-linked allele is greater in females owing to the unique transmission of this chromosome, and therefore the X chromosome is more strongly selecting on the female phenotypic effects of alleles than their male phenotypic effects (Fig. S1). In the case of a dosage-compensated X, on the other hand, the increased selection pressure in males cancels this underlying bias of the X toward female fitness, such that the genetic response—reflecting the phenotypic response—resembles the autosomal case.

The strength of sexually antagonistic selection on individual alleles is often measured by examining male-female *F_ST_*, with the expectation that alleles undergoing positive viability selection in one sex and negative viability selection in the other will exhibit within-generation frequency differences between males and females (Cheng and Kirkpatrick 2016; Ruzicka and Connallon 2022). In line with our observation that phenotypic change is caused by frequency changes at many loci with relatively small effects, we observe low average levels of *F_ST_* at loci underlying the phenotype after the shift in optimum. On autosomes, the average *F_ST_* was 0.00006 (with a mean maximum *F_ST_* across replicates of 0.0008); on the non-dosage compensated X chromosome, the average *F_ST_* was 0.0005 (with a mean maximum across replicates of 0.0010). We should note that, while this suggests that the signature of sexually antagonistic selection on a polygenic phenotype is unlikely to involve high levels of *F_ST_* between the sexes across all loci, or to induce strong *F_ST_* outliers, it is still possible that systematic patterns of male-female *F_ST_* across the genome due to sexually antagonistic selection can be detected (Kasimatis et al. 2021; Ruzicka and Connallon 2022).

#### The effect of dominance

A potential complicating factor for the different responses of the X and autosomes to sexually antagonistic selection is the possibility of alleles with variable dominance. The X chromosome is haploid in males, thereby effectively forcing alleles to be dominant in one sex, in contrast to the autosomes, which can more effectively shelter recessive alleles. We therefore also examined a scenario in which the dominance of alleles affecting the phenotype varied. For simplicity, we assumed that the dominance coefficient of each allele was the same in males and females. Note that here we define dominance in terms of an allele’s effect on the phenotype, not in terms of its effect on fitness, the the usual context in which dominance has been considered in the sexual antagonism literature (Rice 1984; Frank and Patten 2020).

We see very little difference in the relative contributions of dominant vs. recessive alleles to phenotypic change on the X chromosome, compared to the autosomes, despite the fact that recessive alleles should be more ‘visible’ to selection on the X due to their hemizygous expression in haploid males (Fig. S3; Charlesworth et al. 1987). These results can be understood in light of the analyses of Orr and Betancourt (2001), who consider the relative rate of adaptation via beneficial mutations across X-linked and autosomal loci. Their one-locus results demonstrate that the probabilities of fixation of previously-deleterious newly-beneficial alleles are essentially independent of dominance when adaptation occurs from the standing variation. This is because recessive deleterious alleles persist at higher copy number under mutationselection balance than dominant deleterious alleles, but each recessive allele is also less strongly selected when it becomes beneficial (Haldane’s sieve).

By this logic, despite the difference in dominance patterns induced by X-linked vs. autosomal transmission, dominance will not substantially affect the probability of fixation of alleles across the two cases. While we consider an explicitly polygenic phenotype in which fixation of alleles is rare during short-term adaptation, in contrast to Orr and Betancourt’s (2001) single-locus model of selective sweeps from standing variation, the intuition about the lack of impact of dominance in the two models is similar. In our model, beneficial alleles, with phenotypic effects aligned to the shift in optimum, will generally have been selected against due to underdominance induced by stabilizing selection prior to the shift in optimum, and adaptation also primarily occurs via the increase in frequency of alleles from the standing genetic variation (Orr and Betancourt 2001). We observe a negative relationship between the dominance of an allele and its frequency prior to the shift in optimum (Fig. S4), consistent with the view that dominant alleles are at lower frequency and that this trade-off drives the lack of difference in the use of dominant vs. recessive alleles on the X chromosome compared to the autosomes in our model (Fig. S3).

#### Coevolution of the X chromosome and autosomes

Thus far, we have considered the evolutionary dynamics of sex chromosomes and autosomes in isolation of one another. This has generated intuition for the distinct evolutionary responses of the autosomes and the sex chromosomes to sexually antagonistic selection.

In the many taxa with heterogametic sex determination, however, sex chromosomes and autosomes co-exist within the same genome (Bachtrog et al. 2014). We therefore examined the evolutionary dynamics of a genome with both autosomes and sex chromosomes. The details of the dynamics of sexually antagonistic selection in a genome containing both sex chromosomes and autosomes will of course depend on the relative proportions of the genome that they comprise (and other features such as their relative gene content, recombination process, etc.), which vary extensively across taxa. For simplicity, in our analyses, we have chosen to simulate a genome based on the karyotype and recombination patterns of *Drosophila melanogaster*, though allowing autosomal recombination in males and considering both the absence and presence of dosage compensation (see Methods). The results below are, however, are robust to there being no autosomal recombination in males (Fig. S7).

If we consider the fate of a polygenic phenotype encoded across a *Drosophila*-like genome, we find that the overall path to the new sexually antagonistic optimum closely resembles that of an entirely autosomally-encoded phenotype (Fig. 3). This is not unexpected, as even in the case of a *Drosophila*-like genome, with relatively few autosomes and a large X chromosome, the majority of loci underlying the phenotype are nonetheless autosomally encoded, so that the response of autosomal loci dominates the overall phenotypic response to sexually antagonistic selection.

We can also examine the specific contributions of the X chromosome and autosomes to this overall phenotypic response. We initially examine the case of no dosage compensation. In this case, the X chromosome responds very strongly to selection in females, rapidly pushing the phenotype towards the female optimum and not the male optimum (Fig. 3). Countervailing this effect of the X chromosome, the autosomes evolve to contribute more to phenotypic movement to the male optimum than to the female optimum (Fig. 3). These dynamics are consistent with previous theoretical work on the potential for co-evolution between genomic regions in response to sexual antagonism (Wade and Drown 2016; Ågren et al. 2019; Wade and Fogarty 2021).

**Figure 3:**
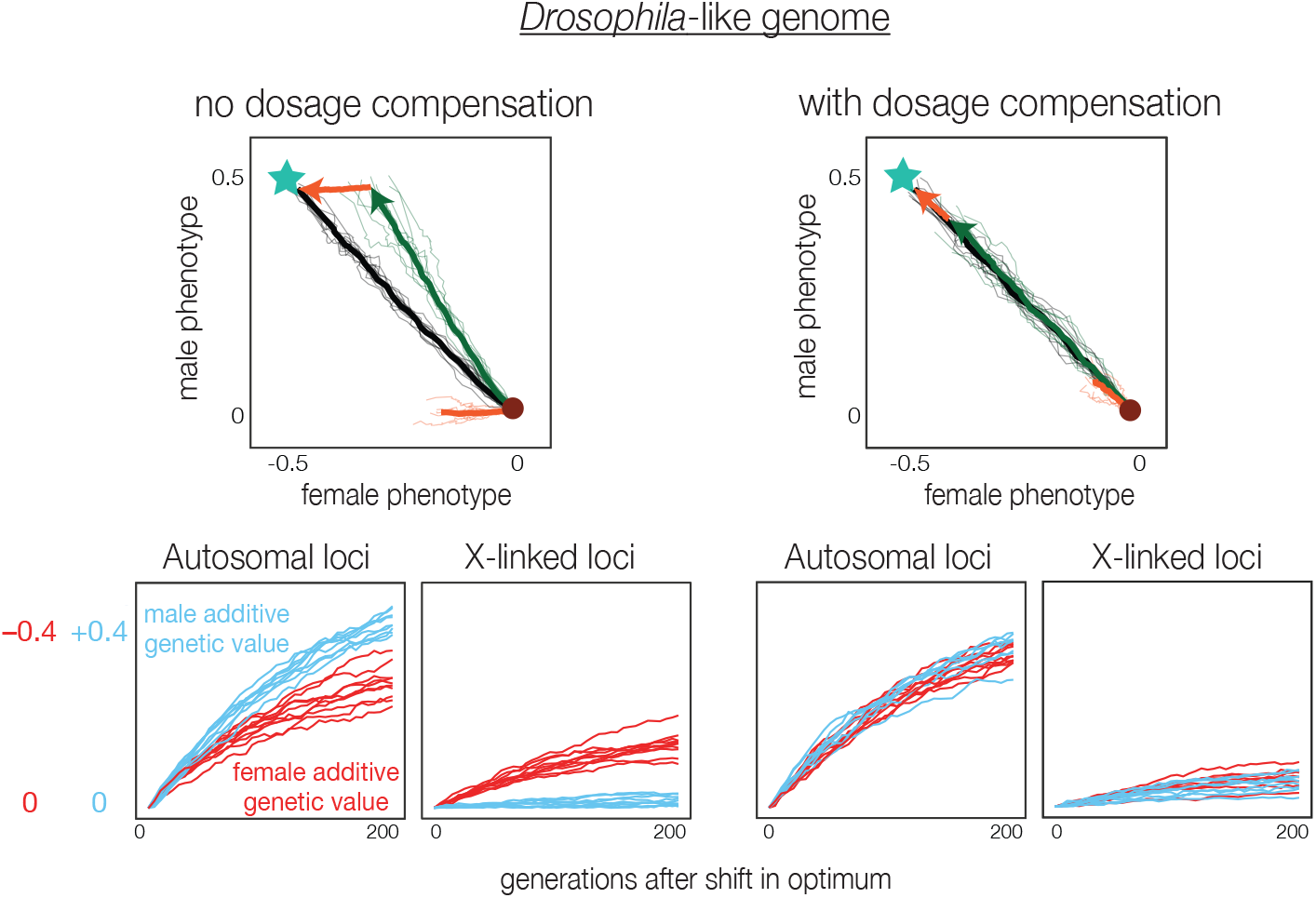
Phenotypic response to sexually antagonistic selection of X-linked and autosomal loci within the same genome. We simulate the response of a phenotype encoded across a *D. melanogaster*-like genome structure, both where the X chromosome does not (left) and does (right) experience dosage compensation. Top panels show the overall path that the mean phenotype takes across the male-female phenotype landscape (black), which is similar in both cases. Under no dosage compensation, however, the X chromosome contributes only to movement toward the female phenotypic optimum (orange path in left panel), while it contributes equally to movement toward the male and female optima when dosage-compensated (orange path in right panel). In the case of no dosage compensation, the autosomes contribute more to movement toward the male phenotypic optimum (green path), to compensate for the X’s female biased effect. Transparent lines show trajectories from individual simulations; bold lines show average trajectories across 10 replicates. To account for drift across chromosomes in their contributions to the trait during the initial burn-in period in our simulations, trajectories are normalized according to their starting values, so that all start at exactly (0,0) in the male-female phenotype space displayed here. Bottom panels track the additive genetic value of autosomes and X chromosomes after the shift in optimum, revealing rapid evolution of a female-biased additive genetic value on the non-dosage-compensated X chromosome and compensatory evolution of a male-biased genetic value on the autosomes.

Essentially, the X chromosome within this genome strongly prioritizes movement to the female optimum, regardless of the effect on male fitness, in the generations immediately following the shift in optimum. In our simulations of an entirely X-linked phenotype, we observed that such a phenotype will eventually respond to selection in males and move toward the male optimum, creating the ‘hockey-stick’ pattern seen in our simulations of an entirely X-chromosomal genome (Fig. 2). However, in the case of a genome containing both a sex chromosome and autosomes, the autosomes provide the predominant contribution to movement toward the male optimum, so that the X chromosome remains largely occupied with movement to the female optimum, as reflected in its strongly female-biased change in additive genetic values (Fig. 3).

**Figure 4:**
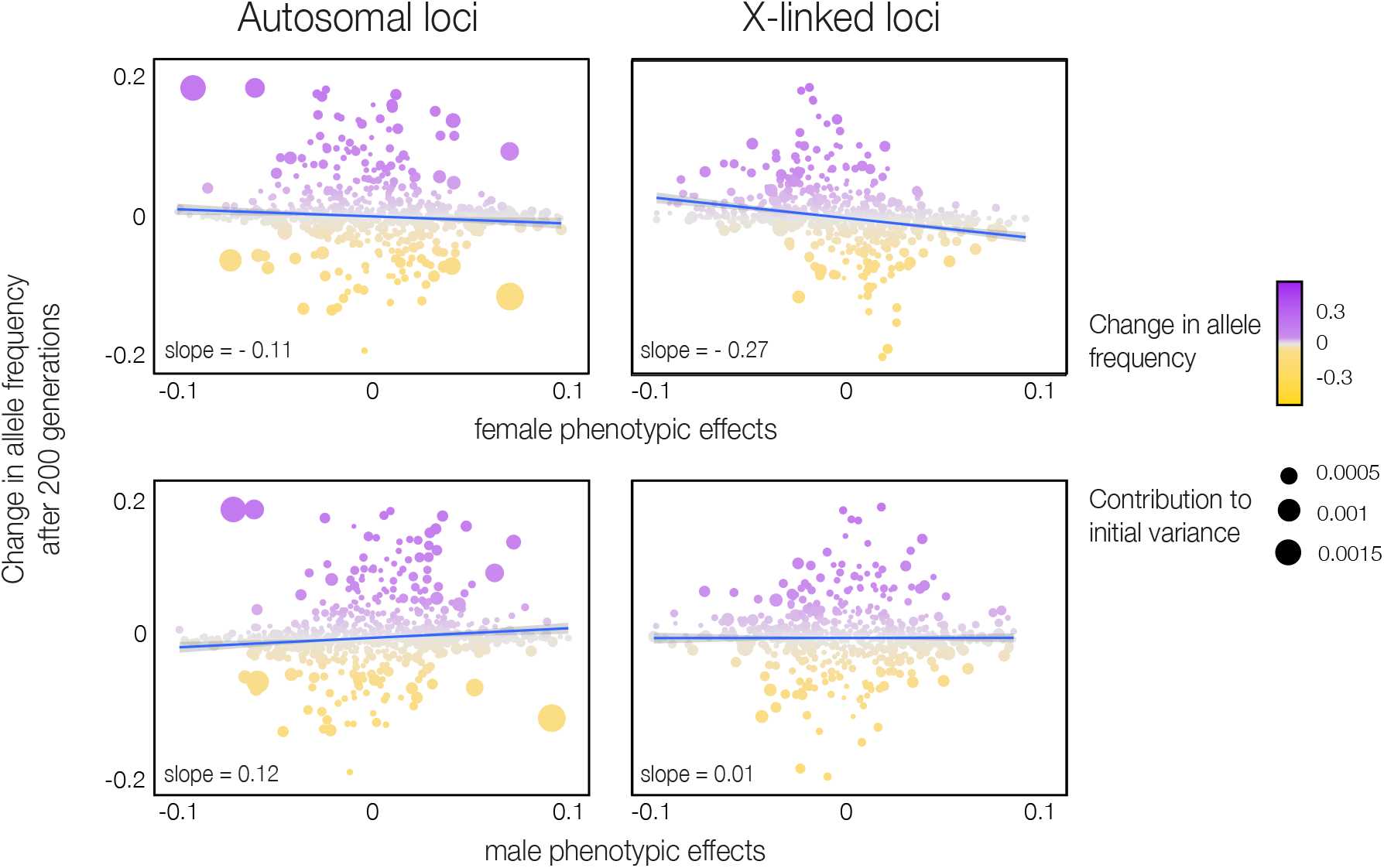
Female-biased genetic response of the X chromosome to sexually antagonistic selection, relative to autosomes within the same genome. Frequency changes of autosomal (left panels) and X-linked (right panels) alleles across 200 generations following a sexually antagonistic shift in male-female optima, plotted as a function of the alleles’ phenotypic effects in females (top panels) and males (bottom panels). Simulations are based on a *D. melanogaster*-like genome without dosage compensation on the X, as in the left panels of Fig. 3. In each case, we randomly sampled 1000 alleles segregating on the X chromosome (right panels) and on the autosomes (left panels) at the time of the shift in optimum. Alleles segregating on the X chromosome show little evidence of selection based on their male phenotypic effects (weak correlation with change in frequency), but show strong evidence of selection based on their female phenotypic effects (strong correlation with change in frequency). In contrast, autosomal alleles show evidence of selection based on both male and female phenotypic effects. Note that, because the new female optimum is lower than the initial optimum, while the new male optimum is higher, selection favors alleles with negative phenotypic effects in females (negative corelation of frequency change with effect size) and positive phenotypic effects in males (positive correlation).

To understand these patterns in more detail, we can examine the frequency change of individual alleles across the X chromosome and autosomes. We find, as expected, that the X chromosome shows a pronounced pattern of selection for alleles with phenotypic effects that move females closer to their optimum, but no similar pattern of selection for male phenotypic effects, while the autosomes show evidence for selection on both male and female phenotypic effects (Fig. 4).

The distinct contribution of X-linked compared to autosomal loci to phenotypic change in males and females in the case of no dosage compensation is unique to sexually antagonistic selection. When the male and female optima shift in a concordant direction, there is a slight difference in the contributions of X-linked loci to phenotypic change in males and females, but this effect is much reduced in magnitude relative to the case of a sexually antagonistic shift in the male and female optima (Fig. S8).

We now consider the case of dosage compensation, with the X up-regulated in males. In this case, we observe a very different pattern. Consistent with our previous simulations of a phenotype encoded entirely on a dosage-compensated X chromosome, the X’s contribution to male and female phenotypic changes is approximately equal in this case (Fig. 3). With no need to balance out an unequal contribution of the X, the autosomes also contribute approximately equally to the male and female phenotypic change. This is also true in the case where the male and female optima shift concordantly (Fig. S8).

These results illustrate the importance of separately examining the contribution of different genomic regions: while the overall phenotypic movement resulting from genome-wide allele frequency shifts looks approximately symmetric with respect to the sexes, the contributions of the autosomes and X chromosome to this phenotypic change can be very distinct. It is furthermore clear from these results that details of the genomic architecture of the phenotype of interest, including the size of the sex chromosome and its enrichment for loci that causally affect the phenotype, will impact how the phenotype evolves in response to sexually antagonistic selection. These results also demonstrate that the presence or absence of dosage compensation can dramatically alter the role played by the X chromosome in sexual antagonism.

The distinct patterns of polygenic adaptation on sex chromosomes vs. autosomes further suggest that sexually antagonistic selection could perhaps be detected via estimation of chromosome-specific contributions to mean genetic values, particularly in model organisms through crossing. It is important to note, however, that it is the change in additive genetic contributions across the X and the autosomes, rather than the absolute value of these scores, which is the characteristic feature of sexual antagonism in our simulations; the X and autosomes may accumulate different genetic contributions even when males and females have the same fitness optimum, as long as their combined effect places the mean phenotype at its optimum (Fig. S9). Studying the dynamics of sexual antagonism through chromosome-specific polygenic scores may therefore be especially amenable to experimental settings, in which the onset and strength of selection in males and females can be controlled and the additive genetic values of different genomic regions measured across time.

### 3.2 Evolutionary dynamics once the new male and female optima have been attained

We have described how the polygenic phenotype evolves towards the new sexually antagonistic fitness optimum. As the male and female phenotypes approach the new optimum, the strength of directional selection decreases, and stabilizing selection comes to dominate (Thornton 2019; Stephan and John 2020; Hayward and Sella 2022). In a sense, the problem of sexual antagonism has then been ‘solved’: males and females have reached their new optima and show sexual dimorphism for the phenotype. Concomitantly, at the genetic level, alleles will no longer show the divergent directional selection in males and females that we associate with sexually antagonistic selection. Instead, the evolutionary dynamics at this stage will reflect the action of stabilizing selection, acting to reducing variance in both the male and female phenotype.

How stabilizing selection affects the dynamics of alleles that underlie both the male and female phenotype is therefore the key question in this phase of sexual antagonism. Stabilizing selection can be thought of as inducing under-dominant selection, such that alleles below 50% frequency are selected against, with a strength proportional to their squared effect size on the phenotype (Lande 1976; Simons et al. 2018). Modifying the standard equations that describe allele frequency change under stabilizing selection (e.g., Simons et al. 2018) to incorporate distinct male and female phenotypes, we find the same pattern of under-dominant selection, in which minor alleles with strong effects on either or both the male or female phenotype are selected against because they increase phenotypic variance (Appendix).

Under stabilizing selection, minor alleles with strong effects on both the male and female phenotype suffer especially in this scenario, as they increase phenotypic variance in both sexes. This can be observed by examining the genetic correlation of male and female phenotypic effects of alleles that segregate at different frequencies. Alleles segregating at low frequencies are likely to have arisen recently, and thus reflect the underlying genetic correlation of new mutations. Minor alleles at moderate or high frequencies have been filtered through the sieve of stabilizing selection in both males and females. These alleles are thus unlikely to have strong effects in both sexes, and therefore should show a reduced genetic correlation between the sexes, relative to those at low frequencies. This is the basis of the idea that the quantitative genetic G-matrix should be shaped by long term selection (Steppan et al. 2002; Jones et al. 2003; Arnold et al. 2008). We confirm this prediction in our simulations, which show the expected decrease in the correlation between male and female effect sizes for minor alleles at higher frequencies (Fig. 5).

**Figure 5:**
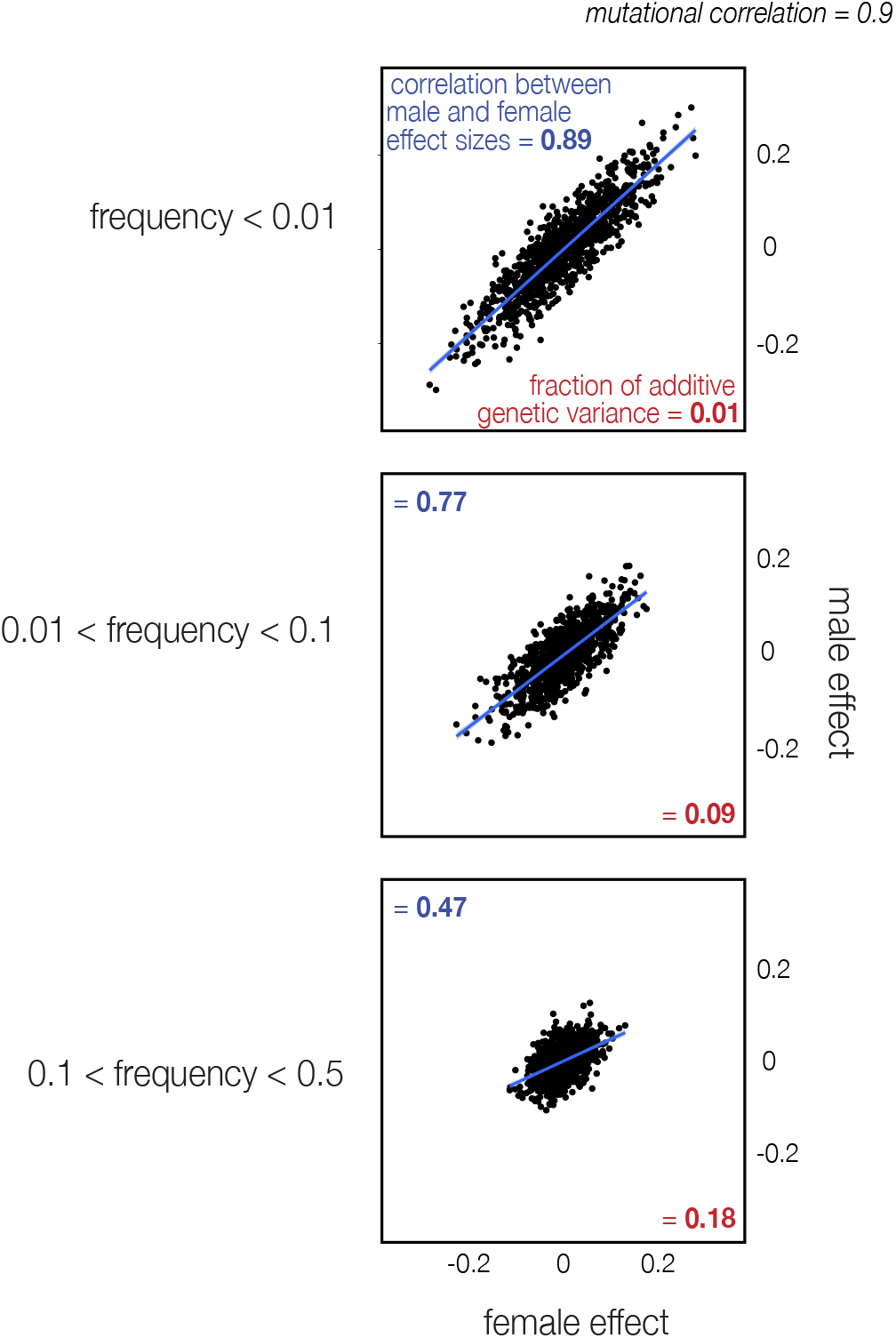
The correlation of the male and female effect sizes of segregating alleles differs across the allele frequency spectrum under stabilizing selection. 5*N* generations after the initial, discordant shift in optimum, we randomly sampled 1000 segregating autosomal alleles from each of three frequency bins, roughly describing minor alleles at low (*p* < 0.01), moderate (0.01 < *p* < 0.1), and high frequencies (0.1 < *p* < 0.5). Alleles in the low-frequency bin show a strong correlation in male-female effect sizes, reflecting an analogous correlation in the underlying mutational distribution. Alleles in the higher frequency bins show a reduced correlation, because stabilizing selection tends to remove alleles with strong effects in both sexes. The samples of 1000 alleles from the moderate and high frequency bins also explain a far greater proportion of the total additive genetic variance (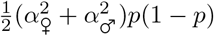 summed across alleles) than the sample of 1000 alleles from the low frequency bin.

These results suggest a novel approach through which the action of stabilizing selection on both males and females could be detected, and its strength potentially estimated: by studying the correlation of male and female effect sizes across the allele frequency spectrum (e.g., using GWAS data). The same phenomenon may also lead to the appearance of a sex-specific genetic architecture, because alleles at higher frequency, which will tend to have more sex-specific effects, explain a greater proportion of the additive genetic variance of a phenotype.

#### The Bulmer effect across the sex chromosomes and autosomes

In species with degraded sex-specific chromosomes, the presence of one copy of the sex-biased X or Z chromosome in the heterogametic sex and two in the homogametic sex will induce differences in the phenotypic variance between the sexes. In addition to these intrinsic effects due to the copy number of the X chromosome, theories such as the ‘unguarded X’ or ‘toxic Y’ predict that the presence of a haploid X or a degenerate Y in males could also result in an increased genetic load in males (Marais et al. 2018; Brown et al. 2020; Xirocostas et al. 2020; Connallon et al. 2022; Sultanova et al. 2023). These effects of the haploid X chromosome on male fitness will naturally also apply in the scenario we have considered.

In the context of stabilizing selection, we identify an additional mechanism through which sex chromosomes can cause differences in genetic load between the sexes: the ‘Bulmer effect’ acting across sex chromosomes and autosomes.

The Bulmer effect describes the negative covariance in phenotypic effects across alleles that is generated under stabilizing selection (Bulmer 1971, 1974). By selecting against extreme phenotypes, stabilizing selection generates negative covariances—or negative linkage disequilibria—between alleles with the same directional effect on the phenotype (i.e., a phenotype-increasing allele is more likely to be transmitted alongside a compensatory phenotype-decreasing allele than alongside another, exacerbatory phenotype-increasing allele). These negative covariances in allelic effects are generated by selection but broken up by recombination in subsequent generations, eventually reaching a stable equilibrium value (assuming the strength of stabilizing selection remains constant).

Previous consideration of the Bulmer effect has focused on the negative covariance generated among autosomal loci (Bulmer 1971, 1974). However, sex chromosomes generate an interesting asymmetry between the sexes in their inherited negative covariance in allelic effects. Consider the case of a species with male heterogamety, but a degenerate Y chromosome. A female in this species will inherit an X chromosome and an autosomal set of chromosomes from her father. This X-autosome set will have experienced stabilizing selection in the same (paternal) genome, generating negative covariance between all loci—autosomal and X-linked—based on their phenotypic effects in males. She will also inherit an X-autosome set from her mother, which will also carry negative covariance genome-wide generated based on alleles’ phenotypic effects in females. In contrast, male offspring inherit only a single X-autosome set, from their mothers, which will carry negative covariance based on alleles’ female phenotypic effects; from their fathers, males receive only autosomes and the negative covariances they contain based on selection on males.

Essentially, females enjoy the benefit of two co-adapted X-autosome dyads, one of which has been ‘tuned’ based on selection in her own sex, while males inherit only one co-adapted X-autosome dyad, which has been tuned based on selection in the opposite sex. Males therefore inherit less negative covariance than females, and therefore exhibit a higher phenotypic variance.

Using the framework of Bulmer (1971), we can calculate the expected equilibrium values for the negative covariance in males and females in this case, and extend these calculations to account for correlated allelic effects between the sexes (Appendix). The relative importance of the Bulmer effect acting between the sex chromosomes and autosomes will depend on details of the sex chromosome system, including the presence or absence of a sex-specific chromosome (Y or W), the size of and recombination rate across the sex chromosomes and autosomes in each sex, and the extent of dosage compensation on the sex-biased chromosomes (X or Z). As an example, we apply our calculations to an organism with a *D. melanogaster*-like chromosome structure (but again with autosomal recombination in males), assuming a focal phenotype under stabilizing selection and controlled by freely-recombining loci evenly dispersed across the X chromosome and autosomes. Even in a *D. melanogaster*-like genome, which has a proportionally large X chromosome, the overall reduction in variance due to negative allelic covariance (the Bulmer effect) is relatively small (around 3% in males and 4% in females) (Fig. 6). Assuming that males and females have equal phenotypic variance prior to the onset of selection, we find that, in equilibrium, the difference between males and females in their inherited covariance is approximately 30% of the total reduction in male phenotypic variance due to the Bulmer effect (Fig. 6). If we relax the assumption of equal initial male and female phenotypic variance to account hemizygosity of the X in males, the difference in the equilibrium inherited covariance between males and females is approximately 20% of the total Bulmer effect in males (Fig. S10).

**Figure 6:**
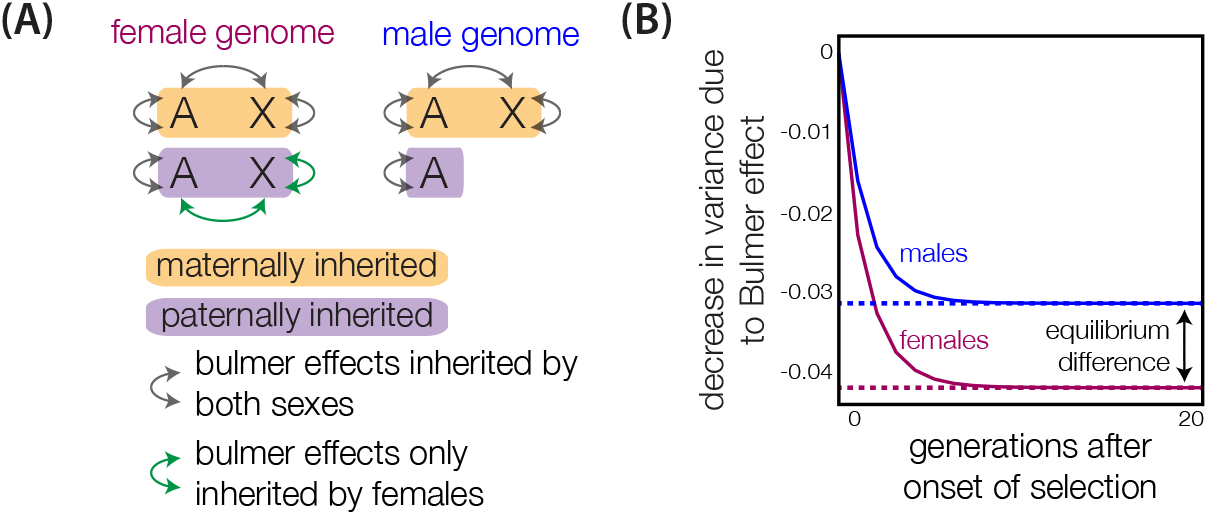
The Bulmer effect across sex chromosomes and autosomes results in higher phenotypic variance in males than in females. (A) Asymmetric inheritance across the genome of the negative covariances induced by stabilizing selection. Because females inherit an X chromosome from their fathers, they inherit negative covariances in phenotypic effects across the X chromosome and autosomes, generated by selection on males in the previous generation. Males, who do not inherit an X chromosome from their fathers, do not inherit these negative X-autosome covariances (see Appendix for a discussion of the case of a non-degraded Y chromosome). (B) In simulations of a *Drosophila*-like genome, the reduction in phenotypic variance due to the Bulmer effect rapidly equilibrates in both sexes, but the overall negative covariance in females is greater than that in males, leaving phenotypic variance higher in males. To isolate this asymmetric impact of the Bulmer effect on males and females, we have assumed that males and females start with the same phenotypic variance at the onset of selection; results for the case in which males have a reduced initial variance owing to their hemizygosity for the X are shown in Fig. S10.

While the absolute reduction in phenotypic variance due to the Bulmer effect is low even in these stylized scenarios, the reductions in males and females are very different because of the asymmetric inheritance of the X chromosome. This should, in theory, result in greater genetic load in males (or in females in ZW/ZZ systems), especially given the ubiquity of stabilizing selection on quantitative phenotypes (Sanjak et al. 2018; Sella and Barton 2019). There is empirical evidence for greater genetic load in the heterogametic sex across taxa (Pipoly et al. 2015; Xirocostas et al. 2020; Sultanova et al. 2023), with this pattern likely due to the confluence of the many mechanisms associated with the inheritance patterns of sex chromosomes (Pipoly et al. 2015; Marais et al. 2018; Brown et al. 2020; Xirocostas et al. 2020; Connallon et al. 2022). The rapidly growing availability of data on sex-determining mechanisms across diverse species offers new opportunities to disentangle these mechanisms in a robust phylogenetic context, and perhaps to characterize the impact of asymmetric Bulmer effects on differences in genetic load between the sexes (Bachtrog et al. 2014; Tree of Sex Consortium 2014).

### 3.3 Maintaining sexually antagonistic selection

Across all the scenarios we have considered, we have observed that a polygenic phenotype under sexually antagonistic selection will rapidly adapt to the new male and female optima, despite a strong male-female genetic correlation. Once at the new optimum, directional selection on the phenotype decreases relative to stabilizing selection. As sexually antagonistic selection can be broadly defined as opposing directional selection on males and females, the signature of sexually antagonistic conflict will therefore also dissipate when the population nears the new optimum. In essence, when the population has reached the new male and female fitness optima, the problem of sexual antagonism has been resolved by the divergence of male and female phenotypic values—that is, the evolution of sexual dimorphism.

However, although we observed rapid resolution of sexual antagonism in our simulations, extensive empirical evidence suggests ongoing sexually antagonistic selection in a wide variety of species (Bonduriansky and Chenoweth 2009; Mank 2017a; Ruzicka et al. 2020). This suggests that the directional phase of the dynamics we have described may be more extended than we have observed in the models presented thus far. We suggest three potential categories of evolutionary constraints that could extend the period of directional selection, and thus increase the probability of detecting ongoing sexually antagonistic selection.

The first category involves limits on mutational input. We have considered a realistic scenario in which alleles underlying the phenotype have correlated, but not identical, phenotypic effects in males and females. If this correlation were stronger, however, movement in an anti-correlated direction towards a new sexually antagonistic optimum would be correspondingly slower (indeed, if the correlation were perfect, with all alleles having identical phenotypic effects in males and females, there could be no anti-correlated movement and therefore no resolution to sexual antagonism).

In our simulations, we have also assumed a particular mutation rate; were the mutation rate lower, and the level of standing variation correspondingly lower, movement to the new male and female optima would also take longer. Similarly, if the average effect of individual mutations is decreased relative to the magnitude of the shift in optimum, movement to the optimum will take longer (Fig. S5).

The second category of constraints that could prolong the directional phase of sexually antagonistic selection involves limitations on selection. If the alleles underlying our focal phenotype, for example, also pleiotropically influenced another phenotype whose male and female optima remained the same, this would exert a considerable restraint on the ability of the focal phenotype to reach a new optimum. New evidence suggests that pleiotropy is a widespread phenomenon for many common phenotypes (Sella and Barton 2019), and even that many phenotypes may in fact be ‘omnigenic’, or genetically controlled by highly interconnected regulatory networks (Boyle et al. 2017). In such scenarios, divergent adaptation of a phenotype to different male and female optima may be prevented by selection on a different phenotype with a shared genetic basis.

The third category of constraints involves changes in environment. We have modeled the evolution of a population to a single new sexually antagonistic optimum. However, in reality, populations experience continual shifts in their adaptive landscape, with male and female fitness optima potentially shifting repeatedly due to environmental changes or inter-specific interactions. In such dynamic adaptive landscapes, a population will continuously chase after different fitness peaks over time.

**Figure 7:**
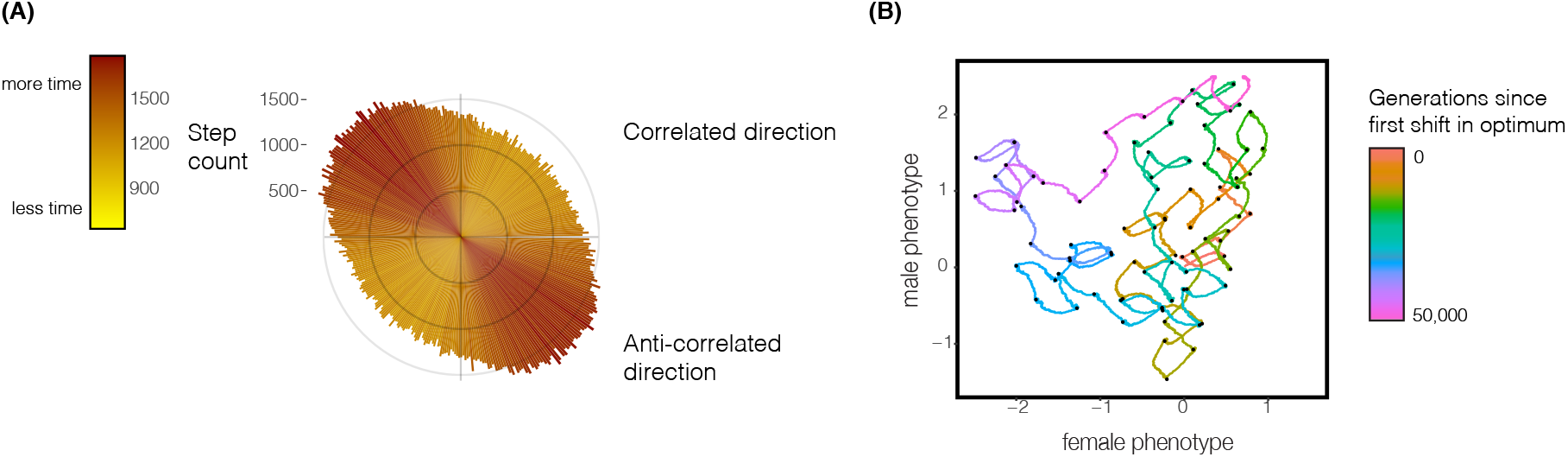
Random fluctuations of the adaptive landscape can generate ongoing sexually antagonistic selection. We simulate a population with a *Drosophila*-like genome structure, as before. The male-female fitness optimum changes every 500 generations, moving a constant distance each time in male-female phenotype space but at a random angle. (A) Histogram of the number of generations in which the population moves in various directions in male-female phenotype space (pooled across replicate simulations of 50, 000 generations). Upward-right (~45°) and downward-left (~225°) movements follow the axis of the underlying male-female genetic correlation of mutations, while downward-right (~135°) and upward-left (~315°) movements are orthogonal to this underlying mutational correlation, and induce sexually antagonistic selection. (B) A representative path taken by the mean phenotype across male-female phenotype space in response to continual shifts in the fitness optimum (black dots).

Such scenarios could result in sustained sexually antagonistic selection. To explore this possibility, we simulated a fluctuating adaptive landscape, in which the male and female fitness optima for a phenotype continually change. In these simulations, the total distance from one optimum to the next in male-female phenotype space is held constant, but the angle at which the jump occurs is random. Under these conditions, we find that, despite the direction of each shift in the male-female optimum being random, the population spends the majority of time evolving in an anti-correlated direction (Fig. 7). This is because phenotypic adaptation in the same direction as the genetic correlation between males and females is very rapid, as standing genetic variation is plentiful along this axis. In contrast, movement in an anticorrelated direction is comparatively slow, because less mutational variation is available along this axis. Consistent with this reasoning, the effect is enhanced if the underlying correlation in male and female phenotypic effects is increased (Fig. S11). Therefore, although the population covers, on average, the same amount of ground in each direction, the duration of time it spends moving in an anti-correlated direction—and thus, experiencing sexually antagonistic selection—is longer. These results suggest that continual random fluctuations in the adaptive landscape alone could induce ongoing sexually antagonistic selection on polygenic phenotypes.

## 4 Discussion

We have developed a model to study the response of polygenic phenotypes to sexually antagonistic selection. We have shown that polygenic phenotypes will rapidly adapt to different male and female fitness optima, even in the face of a strong underlying genetic correlation. Furthermore, we have demonstrated that the sex chromosomes and autosomes can take different paths across phenotype space to these new fitness optima, but that this conclusion is senstive to the details of dosage compensation on the X chromosome. We have then studied the evolutionary dynamics once the polygenic phenotype has reached its new male-female fitness optimum, uncovering a subtle role for the Bulmer effect in generating differences in the equilibrium phenotypic variance of males and females. Finally, we have proposed mechanisms that could prolong the directional phase of sexually antagonistic selection, and thus explain its widespread observance across species. While we have phrased our results in the language of male heterogamety and an X chromosome, our results apply equally well to female heterogamety and a Z chromosome, *mutatis mutandis.* Our conclusions therefore apply to a broad multitude of taxa with diverse sex-determining mechanisms.

### Extensions and limitations of the model

Our analyses have attempted to shed light on the basic dynamics of a polygenic phenotype under sexually antagonistic selection. However, in building our models, we have made assumptions about the evolutionary dynamics at play which may not hold broadly in natural populations or across species.

For example, our assumption of equally strong selection in the two sexes is unrealistic, but serves to clarify the basic dynamics underlying opposing selection in males and females, particularly with respect to sex chromosomes. If selection on the male phenotype were stronger (e.g., due to female-biased mate choice (Janicke et al. 2016)), the underlying patterns we observe across sex chromosomes and autosomes might be obscured: a non-dosage-compensated X might then be equally biased toward male and female interests, obscuring the intrinsic female-biased evolutionary interests of non-dosage-compensated X chromosomes that we (and others) have elucidated. Understanding the separate strengths of selection in each sex is therefore essential for accurate prediction of the phenotypic response to sexually antagonistic selection. However, measuring selection in natural populations is difficult. Therefore, controlled selection experiments in the laboratory may be particularly useful in characterizing these dynamics.

We have also assumed a relatively simple population model, excluding demographic complications such as population structure and non-random mating. These factors are known to have an impact on the dynamics of sexually antagonistic selection, and may enhance or impede the ability of a population to adapt to a new sexually antagonistic optimum (Arnqvist 2011; Flintham et al. 2021; Tazzyman and Abbott 2015; Albert and Otto 2005; Muralidhar 2019). It may be especially interesting to examine how demographic processes arising from resource limitation and competition may intersect with sexual antagonism to promote phenotypic divergence between males and females. Theoretical analyses of the speciation process have found that overcrowding around a phenotypic optimum can drive the evolution of niche specification, by changing the fitness landscape in such a way as to create disruptive selection away from the crowded optimum (Rosenzweig 1978; Dieckmann and Doebeli 1999). This logic has also been applied to the divergence of male and female niches within a species, suggesting a mechanism that would, in our model, endogenously shift the adaptive landscape of males and females (Slatkin 1984; Bolnick and Doebeli 2003; De Lisle 2019). Given the prevalence across taxa of sexually dimorphic niche partitioning, it may be useful to consider how the dynamics of sexually antagonistic selection that we have outlined here can interact with ecological or demographic constraints to promote this form of sexual dimorphism, particularly in light of new experimental evidence for this phenomenon (De Lisle 2023).

We have modeled a phenotype that is completely genetic, with no environmental variance. This assumption has allowed us to gain a clear understanding of the genetic response to sexually antagonistic selection, but, in reality, all polygenic phenotypes will be affected to some degree by an individual’s environment. Environmental noise in the phenotype will likely weaken its adaptive response to to sexually antagonistic selection, making this response more difficult to detect in population genetic data. It is also interesting to consider how different phenotypic responses of the sexes to the influence of the environment may affect these results. If, for example, the male phenotype is more plastic than the female phenotype (e.g., Rohner et al. 2018; Stillwell et al. 2010), and thus more prone to environmental influence, even the autosomes—despite their slight (compensatory to the X) bias toward the male optimum in our wholegenome simulations—could show a more rapid rate of movement toward the female fitness optimum relative to the male optimum. This would simply be due to a difference in how efficiently selection can act on the male vs. female phenotype.

In our analyses of the influence of dosage compensation, we have assumed a mechanism that (1) operates by doubling the effect sizes of alleles on the X chromosome in males (a proxy for up-regulation of the X chromosome in males) and (2) applies uniformly across the X chromosome. With regard to assumption (1), mechanisms of dosage compensation vary widely across species. The system we have described here is found, for example, in *Drosophila melanogaster,* but dosage compensation can also be achieved by down-regulating the X chromosome in females (as in *C. elegans*) or by inactivating a single copy of the X randomly (as in eutherian mammals) or non-randomly (as in marsupials, where the paternal X is inactivated) (Heard and Disteche 2006; Conrad and Akhtar 2012; Mank 2013; Lau et al. 2014; Gu and Walters 2017). These diverse mechanisms of dosage compensation could induce different selection pressures on the X chromosome, and the corresponding response to these selection pressures may depend on the phenotype of interest. For example, the timing of random X inactivation and subsequent development in eutherian mammals could lead certain tissues in females to be effectively hemizygous in their expression of X-linked genes (expressing only the maternal or the paternal X), while others may contain a mosaic of cells, some expressing the maternal X and others the paternal X. Polygenic phenotypes associated with these different tissues may therefore show very different patterns of adaptation on the X chromosome (Wu et al. 2014; Tukiainen et al. 2017; Shvetsova et al. 2019). The tremendous impact of dosage compensation on the dynamics of sexual antagonism in our simulations suggests that explicitly considering the various mechanisms of dosage compensation in these dynamics could be a fruitful avenue for future work.

With regard to assumption (2) above, there is increasing evidence that dosage may not be compensated chromosome-wide in many taxa, with varying in degree along the sex chromosomes (Mank 2013). Our results for dosage compensation vs. no dosage compensation may therefore be thought of as two ends of a continuum, with taxa in which dosage compensation is not complete expected to show intermediate versions of our results.

We have also assumed a particular relationship between the phenotypic effect of an allele and its response to dosage compensation. In our models of the X chromosome without dosage compensation, we assume the phenotypic effect of an allele on the X chromosome in males is 0.5*e_m_*, while in the case with dosage compensation, the phenotypic effect of that allele is *e_m_*; that is, we assume a linear relationship between the phenotypic effect of an allele and the overall expression and copy number of the X chromosome. While there is evidence for such a relationship for polygenic phenotypes in humans (Sidorenko et al. 2019), it is a simplification of the complex array of mutations affecting polygenic phenotypes and their relationship to gene expression; there are also categories of alleles (e.g. alleles containing knockout mutations at protein-coding genes) for which this relationship is unlikely to hold. Future work incorporating more explicit models of gene expression may provide greater insight into precisely how dosage compensation can affect the evolutionary dynamics of sexual antagonism on the sex chromosomes.

Finally, we have assumed that the underlying mutational distribution of male and female effects stays constant throughout the adaptive process. In our model, this mutational distribution represents the extent of the shared underlying genetic architecture of the phenotype, and thus the constraint on the ability of the phenotype to evolve towards separate male and female optima (Lande 1980; Walsh and Blows 2009). The stronger the correlation between male and female effect sizes, the stronger the intragenomic conflict generated by divergence of male and female fitness optima. While we have treated this correlation as a constant parameter, the underlying mutational distribution of effect sizes may itself evolve over time. We predict that, given continual random shifts in the male and female optima as we have simulated above, selection would act to reduce the mutational correlation in male and female effect sizes, allowing the population to adapt more efficiently in any direction. In contrast, if the optimum tends to shift in a correlated direction, i.e., non-randomly, an increase in the underlying mutational correlation would be favored. While evolutionary changes to the distribution of effect sizes are likely to occur on slower time scales than adaptation to new male-female fitness optima, they may be relevant for the long-term impact of sexual antagonism within a species (Arnold et al. 2008; Mank 2017b).

### Sexual antagonism and sex-specific chromosomes

Thus far, we have considered the dynamics of sexually antagonistic selection across only the X (or Z) chromosome and the autosomes. Our exclusion of the sex-specific (Y or W) chromosome from this analysis was because, in many taxa, it is highly degenerate and gene-poor. However, a large Y chromosome may exist in species in which the sex chromosome system has recently evolved or turned over. In such cases, the Y could substantially influence a polygenic phenotype under sexually antagonistic selection. Two possible scenarios are of interest here with respect to the dynamics of sexually antagonistic selection. First, in species with substantial recombination along the lengths of autosomes in males, the young X and Y chromosomes might still recombine along much of their lengths, at least until recombination suppression evolves. That is, the sex chromosomes will harbor a large pseudo-autosomal region, the loci in which behave approximately autosomally in genetic transmission. In such situations, the response of the sex chromosomes to sexually antagonistic selection will resemble that of the autosomes.

Alternatively, in species where males do not recombine autosomally (e.g., *Drosophila*) or restrict crossovers to extreme terminal regions of the chromosomes (e.g., Berset-Brändli et al. 2008), alleles on a newly evolved Y chromosome will not be exchanged with X chromosome alleles, and will therefore experience a male-specific evolutionary trajectory in which their evolutionary ‘interests’ are distinct from those of the autosomes and the X chromosome.

In such a case, the Y chromosome exists only in males and therefore does not experience sexually antagonistic selection per se. However, the Y could still play an important role in sexually antagonistic selection across the genome.

From a genotype-forward perspective, the Y chromosome can act as a safe harbor for sexually antagonistic alleles. If, for example, a male-beneficial female-costly allele translocates from the autosomes or the X chromosome to the Y, this would resolve the sexually antagonistic conflict created by the allele. This is unlikely to be a common mode of resolution for sexually antagonistic selection on a polygenic trait, due to the large number of loci that affect the phenotype genome-wide.

However, a different way in which the Y could shape the dynamics of sexually antagonistic selection on a polygenic phenotype is through its contribution to the evolution of additive genetic values. We would expect Y-linked loci to evolve solely in response to selection in males, and thus rapidly to evolve an additive genetic value that pushes the phenotype toward the male optimum. In our X-autosome simulations, we found that that the autosomes contribute more to movement to the male optimum to compensate for the female bias of the X chromosome’s response (Fig. 3). The presence of a large Y would lessen this pressure on the autosomes to compensate for the X—indeed, the autosomes might even contribute predominantly to movement to the female optimum to compensate for the very strong male bias of the Y.

### Consequences for transitions in sex-determining systems

The view of sexual antagonism that we present here is essentially that of a transient intragenomic conflict. Sexually antagonistic selection drives males and females to evolve divergent phenotypes and, once this sexual dimorphism is achieved, the conflict between the sexes over their shared genome is resolved. This process is rapid, though we have explored mechanisms that might prolong sexually antagonistic conflict.

The apparent simplicity, at the phenotypic level, of the rapid resolution of sexually antagonistic conflict conceals, at the genetic level, complex differences that have been set up between sex chromosomes and autosomes in their evolutionary response to this conflict. Critically, these differences in male-female additive genetic values between sex chromosomes and autosomes will persist even once sexually antagonistic conflict has been resolved, compensating for one another precisely to keep the male and female phenotypes at their respective new optima.

These differences in the additive genetic values of sex chromosomes and autosomes may have significant implications for the stability of the sex chromosome system, and therefore may help us understand the remarkable diversity and lability of sex determining systems across taxa. A major aim in the study of sex chromosome transitions is to identify selective forces which may push a population toward a new sex chromosome system, or constrain a new system from evolving (Bull 1983; Bachtrog et al. 2014; Beukeboom and Perrin 2014). Sexually antagonistic selection in particular is considered to be a key force that can both promote and prevent transitions in sex chromosome systems (Fisher 1931; Rice 1986, 1987; Van Doorn and Kirkpatrick 2007; van Doorn and Kirkpatrick 2010). If, for example, a new male sex-determining mutation appears close to a male-beneficial female-costly allele, this neo-Y chromosome would carry a selective advantage in males, promoting its invasion and a possible transition to a new XX/XY system (Van Doorn and Kirkpatrick 2007; van Doorn and Kirkpatrick 2010). Conversely, if an existing Y chromosome has accumulated many male-beneficial female-costly alleles, any sex chromosome transition that involves the production of phenotypically female XY individuals could be selected against, and a transition in the sex chromosome system would become much less likely (Fisher 1931; Bull 1983; Rice 1987).

In our polygenic model of sexual antagonism, an analogous phenomenon to the presence of sexually antagonistic alleles on a sex chromosome is the bias of a sex chromosome’s additive genetic value towards the phenotypic optimum of one sex or the other (e.g., an X chromosome with an additive genetic value that contributes substantially to the female phenotype reaching its fitness optimum; Fig. 3). This phenomenon could also establish a strong selective force opposing the invasion of new sex chromosome systems which change the inheritance pattern of the original sex chromosomes. For example, it is possible to transition from an XX/XY system to a ZW/ZZ system via the spread of a new dominant female-determining mutation on the ancestral X chromosome, such that this mutated X* chromosome becomes a W chromosome in the new ZW/ZZ system (Bull and Charnov 1977; Bull 1983). However, this transition involves the production of phenotypically female X*Y individuals (Bull and Charnov 1977; Bull 1983). If the population had previously experienced sexually antagonistic selection on a polygenic phenotype, the X chromosome will have accumulated an additive genetic value pushing females toward their new fitness optimum. X*Y females might then suffer a significant fitness cost from having only a single X chromosome (and especially so if the Y is large and carries a substantially male-biased additive genetic value). The reduced fitness of these X*Y females, induced by historical bouts of sexually antagonistic selection, will act as a brake on transitions to a ZW/ZZ system.

It will be interesting to examine how transitions in sex-determining mechanisms play out in an explicitly polygenic context, particularly given recent debate on the role of sexual antagonism in the formation and degradation of sex chromosomes (Lenormand and Roze 2022). More generally, the different patterns of polygenic adaptation we have observed across the sex chromosomes and the autosomes could have implications for crosses between populations, or hybridization events between species, that have experienced different historical selection pressures. If one population has experienced stronger sexually antagonistic selection than the other, hybrids may suffer disproportionately due to mismatched phenotypic contributions from the sex chromosomes. This might provide yet another mechanism—among the many already proposed—for why the sex chromosomes play a particularly important role in hybrid fitness (Orr 1997; Charlesworth et al. 1987; Presgraves 2008; Payseur et al. 2018; Veller et al. 2023).

## Acknowledgments

We are grateful to Carl Veller for comments that improved the manuscript and members of the Coop lab at UC Davis for helpful discussions. PM is supported by a Center for Population Biology postdoctoral fellowship and an NSF postdoctoral fellowship. This work was supported in part by the National Institute of General Medical Sciences of the National Institutes of Health (grant NIH R35 GM136290 to G. Coop).

## Appendix

### S1 Modeling evolution to the new male and female optima using the multivariate breeder’s equation

The response of two correlated phenotypes to selection can be approximated using the multivariate breeder’s equation (Lande 1980; Walsh and Blows 2009). In our case, the two correlated phenotypes are the values of a single phenotype in males and females. Given a ‘G matrix’, **G**, containing the additive genetic variances and covariances of the phenotype in the two sexes, the breeder’s equation for the change in value of the two phenotypes across a single generation is

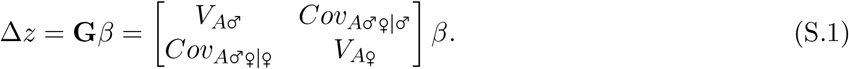

Here, *V*_*A*♂_ and *V*_*A*♀_ are the additive genetic variances of the phenotype in males and females, *Cov*_*A*♂♀|♀_ is the additive genetic covariance between male and female breeding values, calculated in females, and *Cov*_A♂♀|♀_ is the additive genetic covariance between male and female breeding values, calculated in males. 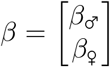 is the selection matrix. The changes in the male and female mean phenotypes can therefore be written separately as

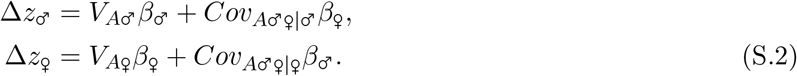

The change in the mean male phenotype reflects a direct phenotypic response due to selection on the male phenotype itself and an indirect phenotypic response due to selection on the female phenotype which, due to the genetic correlation between the male and female phenotype, results in phenotypic change in the male phenotype. The change in the mean female phenotype can be similarly decomposed.

#### An autosomal genome

We first consider the case of an entirely autosomally inherited phenotype. In this case, the covariance terms in the G matrix are equal: *Cov*_*A*♂♀|♀_. Under stabilizing selection on this phenotype toward male and female optimal values *O*_♂_ and *O*_♀_ respectively, the selection matrix is

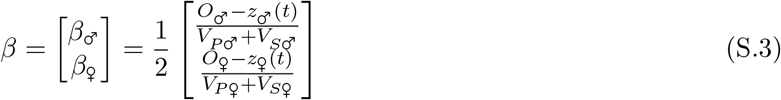

where *z*_♂_(*t*) and *z*_♀_(*t*) are the mean phenotypes in males and females at time *t*, *V*_*P*♂_ and *V*_*P*♀_ are the phenotypic variances in males and females, and *V*_*S*♂_ and *V*_*S*♀_ represent the strength of stabilizing selection on males and females. In all of our simulations, we set *V*_*S*♂_ = *V*_*S*♀_ = 1. The factor of 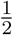 in Eq. (S.3) reflects the fact that we are considering sex-specific selection—because the population is evenly divided into males and females, the expected phenotypic change in either sex is halved relative to that of a phenotype experiencing selection across all members of the population. So, in the case of an autosomally encoded genome, the multivariate breeder’s equation, Eq. (S.1), is

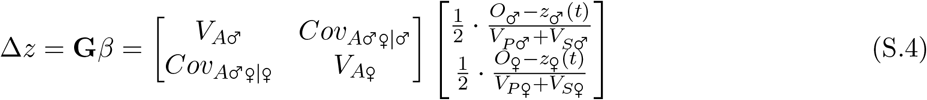

In our simulations of adaptation to the new male-female phenotypic optimum of (+0.5, – 0.5), we can parameterize the multivariate breeder’s equation using the starting estimates of the genetic variances and covariances from our simulations, after a period of 100,000 generations’ burn-in under stabilizing selection around a common optimal phenotype of 0. Note that, because we allow for no environmental effects on the phenotype, *V_A_* = *V_P_* in all scenarios of our model.

We find, as expected under autosomal inheritance, that the starting male and female phenotypic variances are approximately identical (*V*_*P*♂_ = *V*_*P*♀_ = 0.043). We then estimate the starting genetic covariance between the male and female phenotypes by calculating, across all individuals, the covariance between the male and female additive genetic values from their diploid genome (*Cov*_*A*♂♀_ = 0.28). We also calculate, across 10 replicates, the average starting positions of the male and female phenotype, which, as expected, are both approximately 0.

Iterating Eq. (S.4) using these values and plotting the resulting dynamics against those observed in our simulations, we find that the multivariate breeder’s equation provides a very good match to the phenotypic path taken to the new optimum (Fig. 2). We also find that it provides a good approximation to the rate of change of the mean phenotypes in the early generations after the shift in optimum, but that in later generations, the assumption of unchanging additive genetic variance in the breeder’s equation causes discrepancies with the phenotypic rates of change observed in our simulations.

#### An X-linked genome

We now parameterize the multivariate breeder’s equation, Eq. (S.1), for the case of an entirely X-linked genome. We first consider the case in which there is no dosage compensation. There are two ways in which the breeder’s equation must be adjusted in this case, relative to the autosomal case: the selection matrix must be adjusted to reflect the fact that the X spends more time in females than it does in males, and the G matrix must be adjusted to reflect that the X is haploid in males and diploid in females. To account for the former, we adjust the 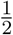 coefficient for selection on males and females in the autosomal selection matrix (Eq. S.3) to 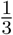 for selection on males and 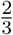 for selection on females:

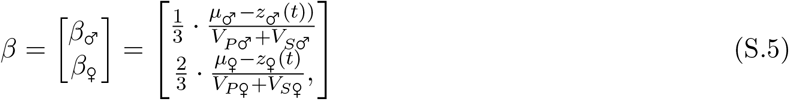

The required adjustment of the G matrix for the X chromosome is more subtle. In the autosomal case, the covariance terms in the G matrix represent the covariance in additive genetic values between the male and female phenotype. However, in the case of the X chromosome, haploidy in males and diploidy in females means that these covariances will differ between males and females, relative to the autosomal case. To account for this discrepancy, we calculate *Cov*_*A*♂♀|♂_ (in the first row of the G matrix) by calculating the covariance of the additive male and female genetic values across all (haploid) male genomes. Similarly *Cov*_*A*♂♀|♂_ (in the second row of the G matrix) is calculated similarly by taking the covariance of the additive male and female genetic values across all (diploid) female genomes (this is the same calculation as for the autosomal case). The average values of these two covariances at the end of the burn-in period of our simulations were, respectively, 0.015 and 0.03, reflecting the doubled additive genetic values in females relative to males due to their two copies of the X chromosome, and consistent with results from previous analyses of the additive genetic variance of X-linked loci (Lande 1980; Fernando and Grossman 1990; Kent Jr et al. 2005; Yang et al. 2011).

In the case of dosage compensation on the X, we add a factor of 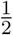 to the additive male-female genetic covariance in males, *Cov*_*A*♂♀|♂_, to reflect the decreased efficiency of selection on additive genetic values in females when ‘translated’ to males. Similarly, we add a factor of 2 to the additive male-female genetic covariance in females, *Cov*_*A*♂♀|♂_, to reflect the inflation of selection on additive genetic values in males relative to females.

With these adaptations to the multivariate breeder’s equation, we observe that it very accurately predicts the path taken to the new phenotypic optimum in our simulations in the case of an X-linked genome with or without dosage compensation (Fig. 2). This suggests that the initial evolution of sexual dimorphism in a polygenic phenotype will often accurately be predicted by the multivariate breeder’s equation.

### S2 Allelic dynamics at the new optimum

Once the male and female phenotypes have reached their new optima, stabilizing selection will prevail around these optima. We adapt the basic model of stabilizing selection (making use of the notation of Simons et al. 2018) to explicitly describe the case of a phenotype under stabilizing selection for different optima in males and females.

In the classical model of stabilizing selection around a *single* fitness optimum, we may consider the case of a particular di-allelic locus, at which the ‘focal’ allele has effect *a* on the phenotype and segregates at frequency *q*, with *p* the frequency of the alternative allele at the locus. We assume that the phenotype is normally distributed with variance *σ*^2^, and experiences weak stabilizing selection, implemented as a Gaussian fitness function with variance *w*^2^. *r* is the displacement of the mean phenotype from its fitness optimum at some point in time. The fitnesses of the three possible genotypes at this locus are then (from Simons et al. 2018):

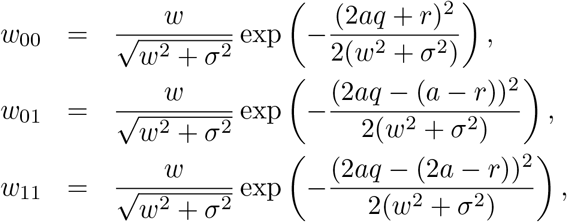

where the subscripts indicate the number of focal alleles carried on each copy of the diploid genome. From these fitnesses, the expected change in the frequency of the focal allele is

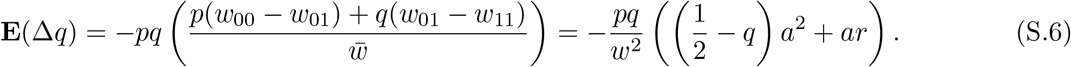

Eq. (S.6) can be broken down into two parts. The first represents stabilizing selection acting on the allele, which is proportional to the phenotypic variance generated by that allele (proportional to *a*^2^). The second represents directional selection acting on the allele, which is proportional to the distance *r* of the mean phenotype from its fitness optimum.

Let us now assume that the focal allele affects a phenotype for which males and females have different fitness optima. We assume that the allele has effects *a*_♂_ and *a*_♀_ on the male and female phenotypic values. We further assume that the the fitness optima for males and females are not extremely distant. The fitnesses of the three genotypes at the focal locus under this model would now be

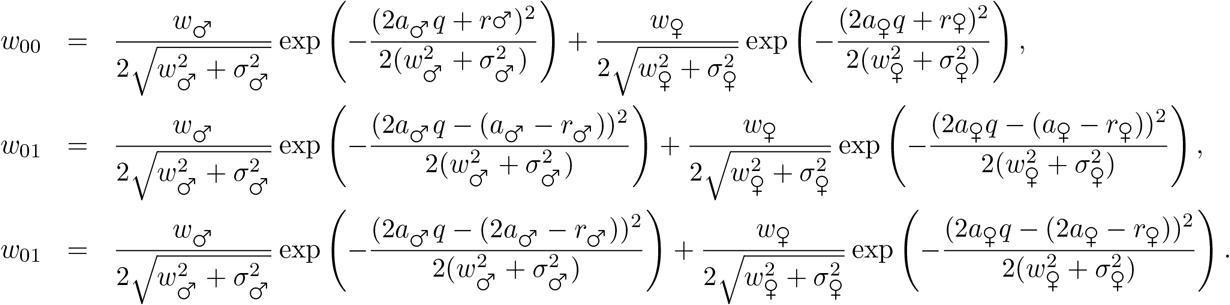

These can be simplified if we assume that the strength of stabilizing selection is equal in the two sexes (*w*_♂_ = *w*_♀_), which further implies that the phenotypic variances of the two sexes are the same (*σ*_♂_ = *σ*_♀_):

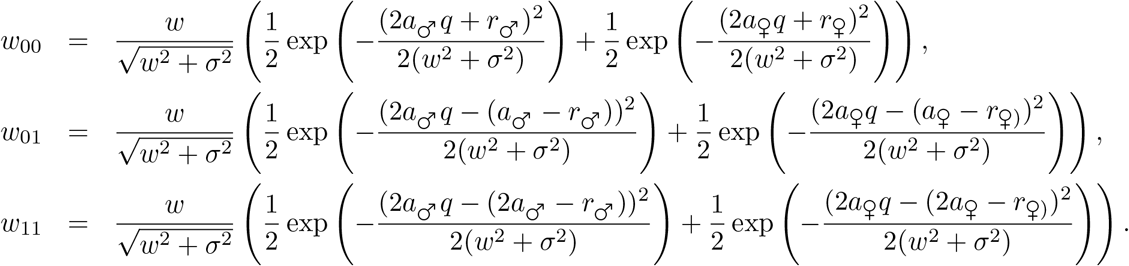

Making the assumption that the Gaussian fitness function is much wider than the phenotypic distribution (*w* ≫ *σ*), as expected under weak stabilizing selection, the change in frequency of an allele under dual stabilizing selection around the male and female optima is

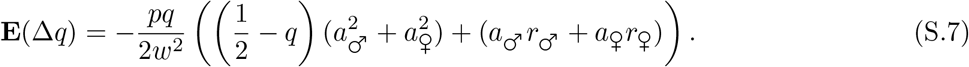

Eq. (S.7) is analogous to the simple case of stabilizing selection of a single phenotype around an optimal value (Eq. S.6), but the change in the frequency of the allele now depends on stabilizing and directional selection in both males and females. When the population is at the male-female optimum, *r*_♂_ = *r*_♀_ = 0, and so directional selection no longer operates, and selection is against rare alleles (*q* < 1/2) without regard to the direction of their effects on the phenotype:

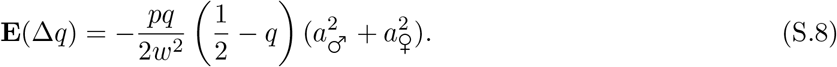

In this case, it is clear from Eq. (S.8) that alleles with strong effects on the phenotype in both sexes (that is, alleles with large values of both 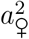 and 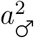) will be especially strongly selected against, relative to alleles with strong effects in only one sex or alleles with weak effects in both sexes. This observation underlies the decrease in the between-sex correlation of allelic effects as we move up the allele frequency spectrum, as illustrated in Fig. S11.

In the case of selection on separate male and female phenotypes, we can also consider what the allelic dynamics above would look like in the case of an X-linked locus. Modifying Eq. (S.6) to account for the transmission of an X chromosome, we find that the expected change in frequency of an X-linked allele is

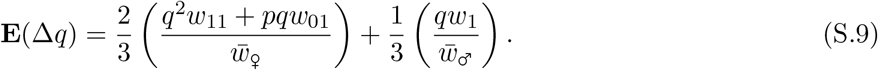

In the case of no dosage compensation, this translates to

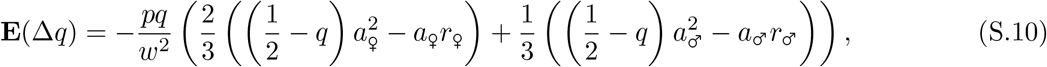

reflecting the increased selection on female, rather than male, fitness due to the asymmetric inheritance pattern of the X chromosome.

The calculations above assume *w* ≫ *σ*, which holds in the case of weak stabilizing selection around a single phenotypic optimum, but they might break down in the case of sex-specific optima if the male and female fitness optima are very far apart so that the variance of the combined male-female phenotypic distribution is large relative to the strength of stabilizing selection (Felsenstein 1979).

### S3 The Bulmer effect across the sex chromosomes and autosomes

Here, we study how the Bulmer effect—the negative allelic covariances generated by stabilizing selection— is be affected by the sex-biased transmission of the X chromosome. In the infinitesimal model of a phenotype under stabilizing selection that Bulmer (1971) studied, the frequencies of individual alleles do not change, and so the reduction in phenotypic variance caused by stabilizing selection is achieved solely by the build-up of negative covariances (i.e., negative linkage disequilibria) among alleles with the same directional effect on the phenotype. Here we modify the basic autosomal model studied by Bulmer (1971) to accommodate an X chromosome, considering a male-heterogametic species with a degenerate Y chromosome (we discuss the potential impact of a non-degenerate Y chromosome at the end of this section).

Adopting the notation of Bulmer (1971) and assuming no environmental effects on the phenotype, an individual’s phenotypic value, *G*, can be described as the sum of the effects of all alleles in their genome. The variation of *G* across individuals then has two components: the genic variance, which describes the variance from alleles segregating at individual loci, and a covariance term from linkage disequilibria across loci:

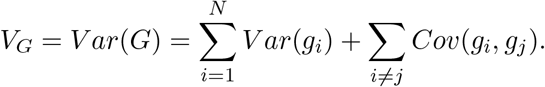

In the case of a genome with both autosomes and an X chromosome, we can partition this overall genetic variance into distinct components deriving from autosomally-encoded and X-encoded loci:

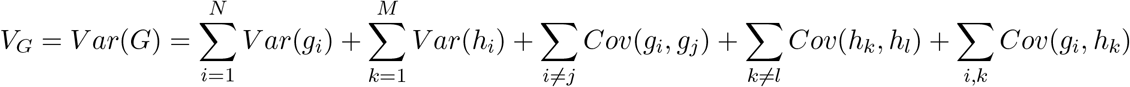

The total genetic variance is composed of the genic variance contributed by the autosomes (*Var*(*g_i_*)) and the genic variance contributed by the X chromosomes (*Var*(*h_i_*)), as well as three types of covariances: the covariances amongst the *N* autosomal loci (∑_*i*≠*j*_ *Cov*(*g_i_, g_j_*), the covariances amongst the *M* X-linked loci (∑_*k*≠*l*_ *Cov*(*h_i_, h_l_*)), and the covariances between autosomal and X-linked loci ∑_*i,k*_ *Cov*(*g_i_, h_k_*)).

Under stabilizing selection, allelic covariances across the genome are expected to be negative. We can calculate how this negative covariance is generated across a single generation of stabilizing selection. We assume that there has been no previous selection on the phenotype and that all loci begin in linkage equilibrium, such that the phenotypic variance is initialy

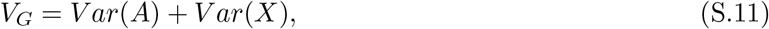

where *Var*(*A*) and *Var*(*X*) equal, in this case, the genic variances contributed by autosomes and the X chromosome. After selection has acted, but before the creation of the next generation, the new genetic variance can be written, following the notation of Bulmer (1971), as

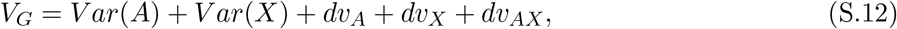

where *dv* represents the negative covariances that stabilizing selection has generated amongst the various sets of loci set out above.

We can then track how this negative covariance is transmitted by males and females to the next generation. For simplicity, we assume that all loci are unlinked, though see Bulmer (1974) for the appropriate modification for variable recombination rates. We also initially assume that allelic effects are identical in males and females.

We first calculate how the negative covariance generated by selection in females, *dv*_≠_, is transmitted as a covariance inherited by their eggs, *di_egg_*:

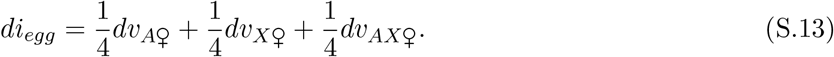

The covariance inherited by eggs is smaller than that generated in females because of (1) recombination, which breaks up the allelic associations in oogenesis, and (2) the presence of only a haploid set of chromosomes in the egg relative to a diploid set in the mother.

We can perform a similar calculation for sperm. Males produce two types of sperm, X-bearing and Y-bearing, which we consider separately. The covariance inherited by X-bearing sperm is

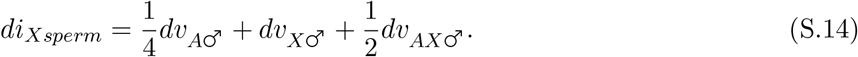

While the first term, describing the dilution of autosomal covariance, is the same as calculated for female gametes, the covariance generated among X-linked loci in males is transmitted undiluted to their sperm because the X does not recombine in males.

Y-bearing sperm, on the other hand, do not inherit any covariance involving the X chromosome, so that

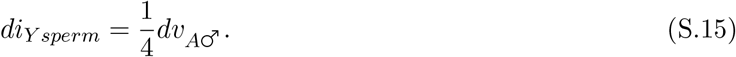

From these equations, we can then calculate the expected covariance inherited by daughters (who inherit covariance via an egg from their mother and an X-bearing sperm from their father):

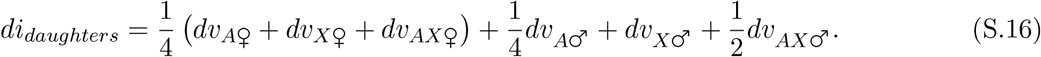

We can also perform the analogous calculation for sons (who inherit covariance via an egg from their mother and an Y-bearing sperm from their father):

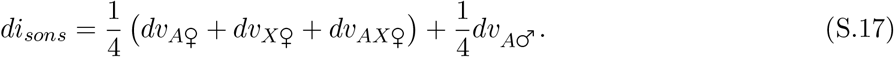

Thus, the expected difference in negative covariance between males and females at the start of the next generation is

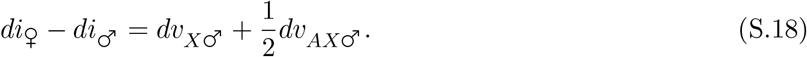

This difference derives from the fact that daughters, but not sons, inherit from their fathers (1) an undiluted X chromosome and (2) a co-adapted X-autosome dyad. Both have accumulated negative covariance due to stabilizing selection in males in the previous generation. The relative importance of these two contributions will depend on the proportion of loci affecting the trait that are X-linked. However, in the realistic case in which the X carries substantially fewer trait-affecting loci than the autosomes, effect (2) above, reflecting the negative covariances between all autosomal loci and all X-linked loci, is likely to be much larger than effect (1), which involves only the negative covariances amongst X-linked loci.

Note that if we considered only autosomal loci, the inherited covariance by both males and females at the start of the next generation would be 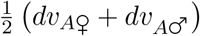. Assuming that selection acts equally in the two sexes, this becomes 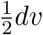, which recovers the original result from Bulmer (1971).

Bulmer (1971) also calculated the equilibrium value of negative covariance, *d*\*, at which the new covariance generated by selection each generation, *d_v_*, is perfectly balanced by the breakdown of inherited covariance *d_i_*. In his model, this is the equilibrium amount by which the phenotypic variance is reduced by stabilizing selection. We calculate this equilibrium for the case of a genome with autosomes and an X chromosome. The update equations for the overall negative covariances in different genomic regions each generation are

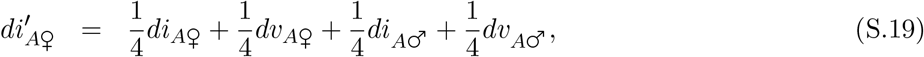

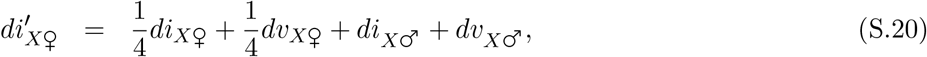

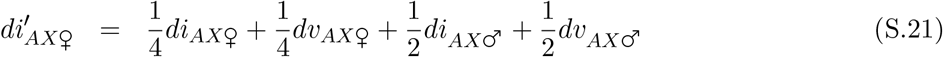

in females and

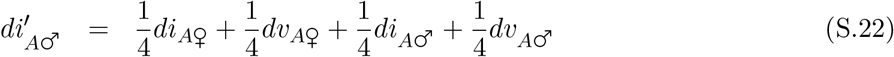

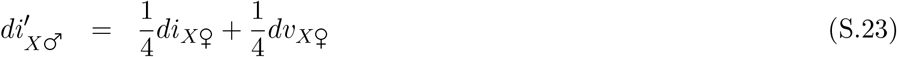

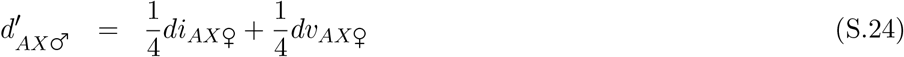

in males.

In equilibrium, the overall negative covariance must be equal in each genomic region; we therefore solve for the equilibrium separately for each genomic region, and then sum these values to find the genome-wide value. Performing this calculation, we find that the equilibrium difference between males and females in their overall negative covariances is

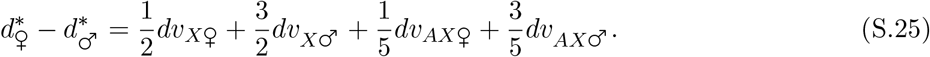

Thus far, we have assumed that allelic effects are identical in males and females. In the results presented in the main text, however, we make the more realistic assumption that allelic effects will be correlated between males and females, but not identical. Under this assumption, the negative covariance inherited by an offspring of the opposite sex will be diluted by a factor *ρ*^2^, where *ρ* is the correlation between male and female effect sizes, so that Eq. (S.25) becomes

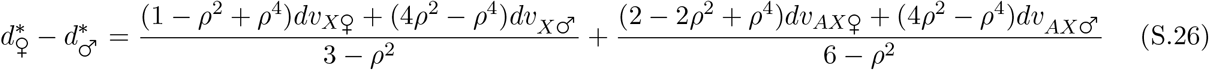

. This simplifies to Eq. (S.25) when *ρ* = 1.

Surprisingly, the consequence of correlated, rather than equal, effect sizes in males and females is to diminish, rather than to amplify, the difference in inherited covariance between the sexes. The difference between males and females in inherited covariance is due to the fact females inherit within-X and X-autosome negative covariances from their fathers, while males do not. If alleles have correlated, rather than identical, effect sizes, males will receive a diluted within-X and X-autosome covariance from their mothers, while females will receive a diluted within-X and X-autosome covariance from their fathers (because covariances generated by selection in one sex become smaller when ‘redeemed in the currency’ of the effect sizes of the other sex). This could, in theory, either enhance or reduce the difference in inherited covariance between the sexes.

However, assuming an equal strength of stabilizing selection in each sex, the transmitted covariance from fathers to daughters on the X and across the X and the autosomes is greater than that from mothers for the same chromosomes because the X does not recombine in males. As a result, the between-sex ‘dilution’ due to correlated effect sizes decreases the covariance inherited by females more than that inherited by males, and therefore reduces the difference in inherited covariance between the sexes.

It is also important to note in these analyses that, following Bulmer (1971), we have remained agnostic about the amount of covariance generated by stabilizing selection in different genomic regions (*dv*). Depending on the genomic distribution of loci underlying a phenotype and the associated phenotypic effects at these loci, *dv_X_* and *dv_AX_* may vary widely and could change the overall equilibrium balance between males and females.

We can use these equations to approximate the expected difference in phenotypic variance between males and females due to the Bulmer effect within a realistic genome. We examine a genome based on the chromosome structure of *Drosophila melanogaster*, although, as in the main text, we allow autosomal recombination in males and assume no dosage compensation (Methods). In such a genome, the X chromosome comprises approximately 17% of the total physical genome. If we further assume that loci are evenly distributed across genomic regions according to their relative physical sizes, then approximately 3% of pairs of loci both lie on the X chromosome, 67% both lie on autosomes, and 29% of locus pairs span X chromosome and the autosomes.

We estimate the strength of stabilizing selection using equation 22 in Bulmer (1971), which states that the decrease in variance due to stabilizing selection in a given generation is 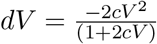, where *V* is the phenotypic variance and 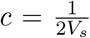, with 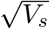 the width of the stabilizing selection function. We assume here that phenotypic effects of alleles are equal in males and females, so that *ρ* = 1.

To study the change in variance between males and females due to the Bulmer effect alone (rather than including the difference due to the haploidy of the X in males), we assume that the starting variance in males and females is equal *V*_♂0_ = *V*_♀0_ = 0.04). We set the width of the stabilizing selection fitness function at 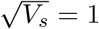 to reflect the parameter settings used in our previous simulations. In this scenario, the Bulmer effect between the X and the autosomes will result in only a 1% difference between the phenotypic variance of males and females. However, because the decrease in phenotypic variance due to the Bulmer effect is only 3% in males and 4% in females, the difference between males and females due to the asymmetric inheritance of Bulmer effects on the X chromosome is actually 30% of the total reduction in males. While the overall reduction in phenotypic variance due to the Bulmer effect is not strong for the parameters we have chosen, the relative difference between males and females in their inherited covariance may be substantial.

So far, we have assumed that the starting phenotypic variance of males and females is equal. However, as males have only one X chromosome and females have two, we expect males to have a reduced phenotypic variance prior to the start of selection (assuming no dosage compensation). If we adjust our calculations so that the starting phenotypic variance in males is decreased proportionally to the number of loci on a single X chromosome, the difference between males and females due to the asymmetric inheritance of Bulmer effects on the X chromosome is approximately 20% of the total reduction in males.

In this example, we have considered a genome with a large X chromosome and a phenotype in which alleles have the same effect sizes in males and females. The relatively small reduction in phenotypic variance that we calculate under these permissive conditions suggests that the the absolute difference in the phenotypic variance of males and females due to the Bulmer effect across the X and autosomes is unlikely to be very large for most taxa. However, depending on the distribution of loci underlying the phenotype, this assymetric Bulmer effect will contribute to an overall higher genetic load in males in combination with other effects, such as the unguarded X or toxic Y (Marais et al. 2018; Brown et al. 2020; Xirocostas et al. 2020; Connallon et al. 2022). All of our results can also be applied to a ZW/ZZ sex chromosome system, albeit with the sexes reversed so that females will have a greater phenotypic variance than males as a result of the transmission of a Z and Z-autosome pair from mothers to sons, and thus suffer a greater genetic load.

In these calculations, we have not considered the impact of a sex-specific chromosome (a Y or W). The presence of a sex-specific chromosome might change our results in two ways, which we will present in the context of an XX/XY system. If a gene-rich Y chromosome is present, but does not recombine with the X in males (as might be a case for a newly evolved Y in a system with male achiasmy), then, because it is transmitted from fathers to sons directly and wholly intact, sons would inherit negative covariance on the Y and from a Y-autosome pair from their fathers. This would reduce the difference in inherited covariance between males and females, and the overall equilibrium difference in phenotypic variance between the sexes. If however, the Y recombines with the X chromosome along much of its length (as expected in the case of a newly evolved sex chromosome pair in a species with recombination in the heterogametic sex), the sex chromosomes will essentially behave like a pair of autosomes and there will be little difference between the equilibrium variance observed in males and females. While this has the same outcome as the previous scenario, it is not because of the presence of a male-specific chromosome, but rather because there is no real sex bias in the transmission of the sex chromosomes.

Generally, the results in this section rely on the assumption of an asymmetry in gene content between the sex-biased (X or Z) and sex-specific (Y or W) chromosomes, which is expected to evolve across the life span of a sex chromosome pair (Bachtrog et al. 2014).

