## Supplementary Figures for "Polygenic outcomes of sexually antagonistic selection"

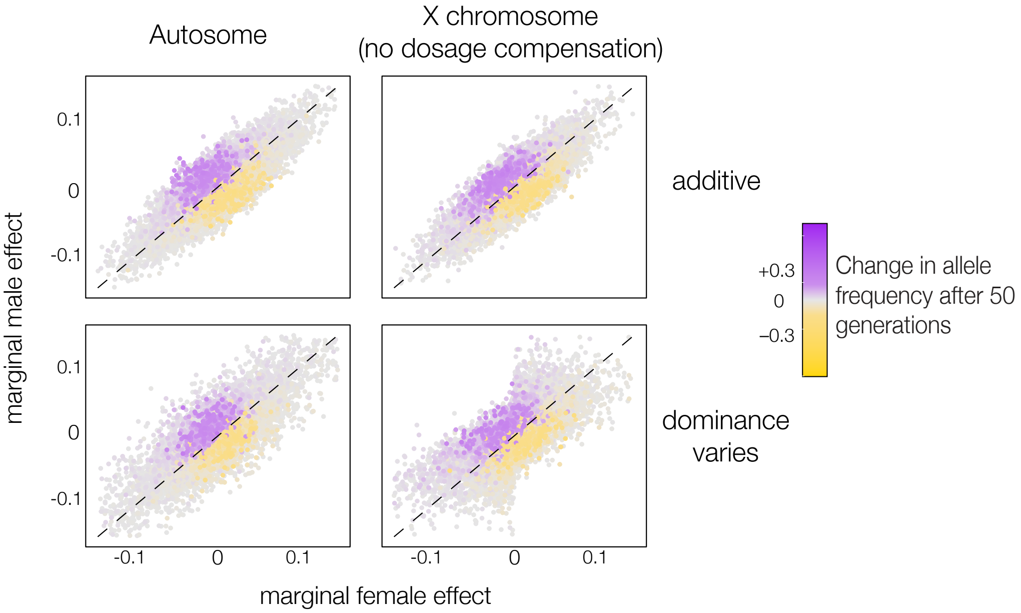


**Figure S1:** As in the right panels of Fig. 2, these panels show changes in frequency across 50 generations for all alleles that were segregating at the time of the optimum shift (color of bubbles). However, rather than displaying this change in frequency as a function of alleles' homozygous phenotypic effects as in Fig. 2, here we show this change as a function of their marginal phenotypic effects in females (x-axis) and males (y-axis). For autosomal loci, we calculate the marginal effect of an allele in females as $(1-p)(h)(\alpha_{♀})+p\alpha_{♀}$ and in males as $(1-p)(h)(\alpha_{♂})+p\alpha_{♂}$. In the case of an X chromosome without dosage compensation, the marginal effect of an allele in females is calculated as in the autosomal case, but the marginal effect of an allele in males is instead $0.5p\alpha_{♂}$, reflecting the haploidy of the X chromosome in males. Because these calculations of the marginal phenotypic effects of alleles take into account the female-biased transmission pattern of the X, we do not observe the female-biased interests of the X chromosome in allele frequency changes as we do in Fig. 2. Note that here, unlike in Fig. 2, all bubbles are the same size.

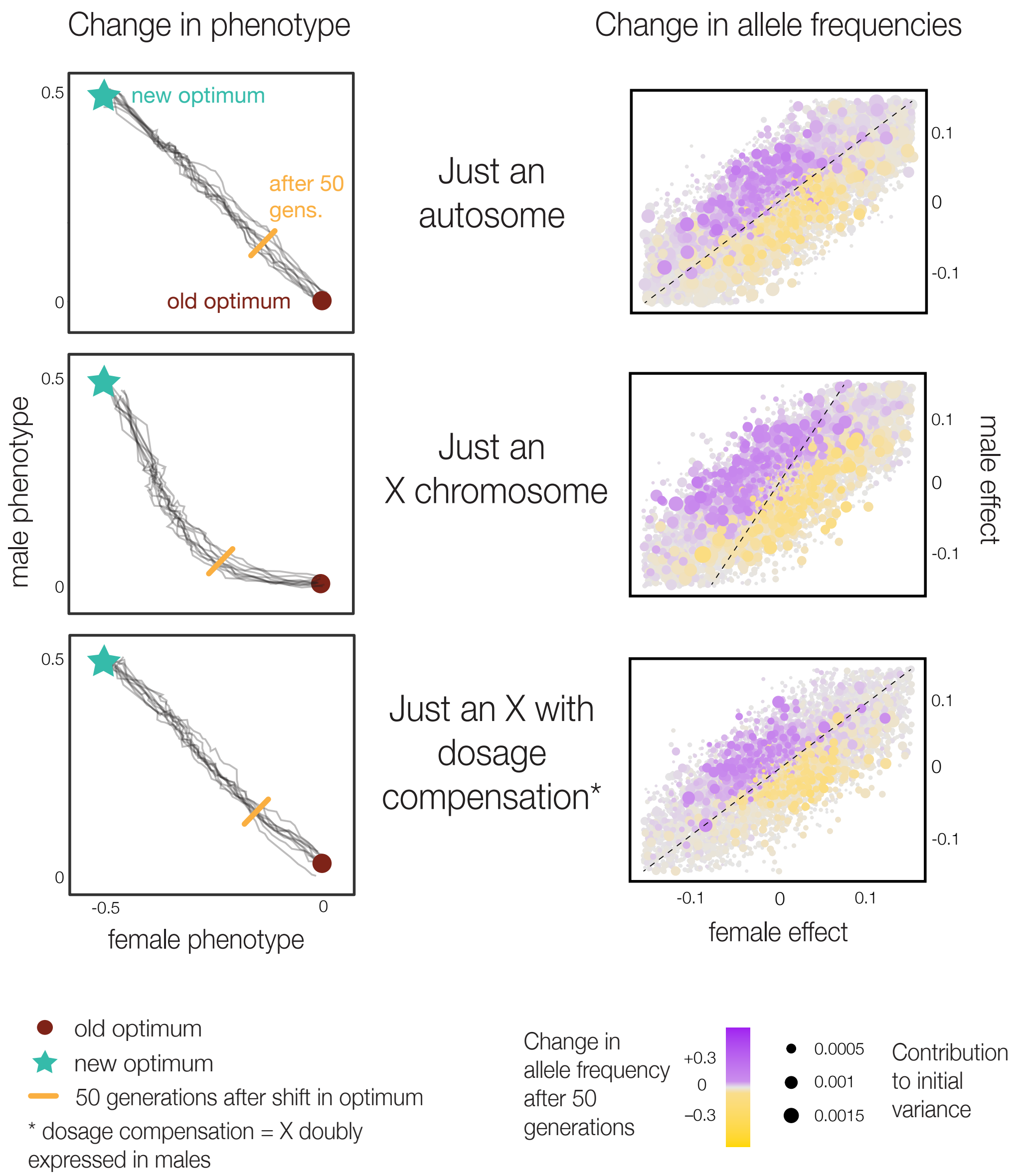


**Figure S2:** Same as Fig. 2, but alleles have variable dominance coefficients. The dominance coefficient of each allele, $h$, is drawn from a from a uniform distribution between 0 and 1, and the effect of an single allele is $he_m$ and $he_f$ in males and females respectively for autosomal loci. The effect of an single X-linked allele is $he_f$ in females, and either $0.5e_m$ or $e_m$ in males depending on whether or not dosage compensation is in effect.

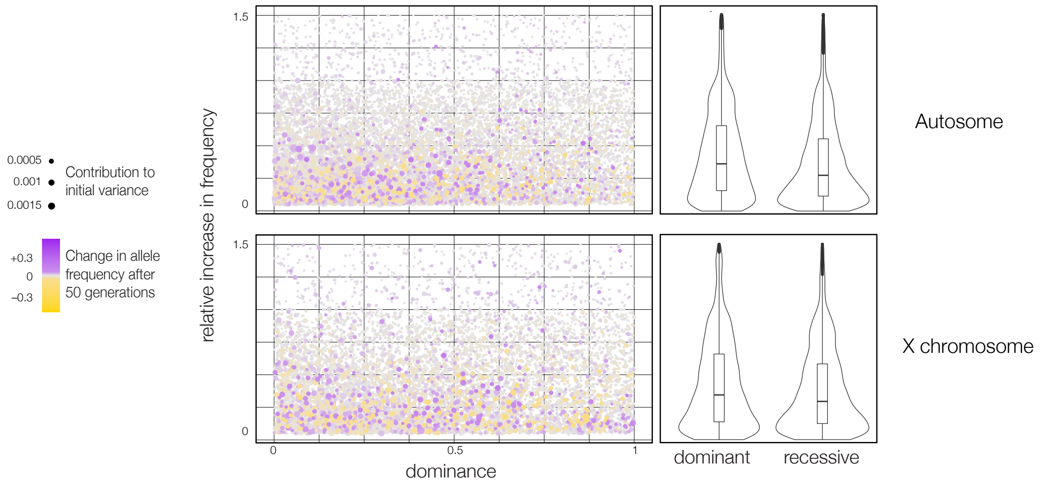


**Figure S3:** The dominance coefficient of an allele does not impact the change in frequency of that allele after a shift in fitness optimum, regardless of whether that allele is X-linked or autosomally. We examine the relative change in frequency after 50 generations for alleles segregating in the population at the time of the optimum shift (the relative change in frequency of an allele is calculated as $p_{50}/p_0$ where $p_t$ is the allele's frequency in generation $t$ after the optimum shift). We see no correlation between the relative change in frequency of an allele and its dominance coefficient ($h$). On the left, we show this relationship across all dominance values ($0 < h < 1$), while on the right, we classify alleles as either dominant ($h >= 0.5$) or recessive ($h < 0.5$). We do not observe a difference between the X chromosome and autosomes in the relative change in frequency of dominant or recessive alleles using either classification.

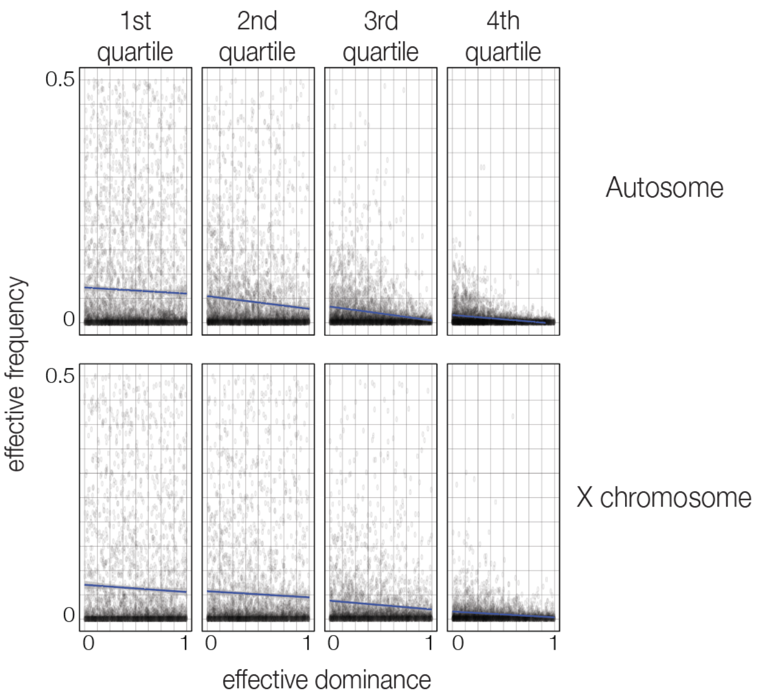


**Figure S4:** There is a negative relationship between the dominance coefficient of an allele and its frequency in the population. Here, rather than considering only derived alleles (see Methods), we examine both derived and ancestral alleles at every locus to capture the full dominance-frequency spectrum. The frequency of these ancestral alleles is simply $1 - p$ where $p$ is the frequency of the derived allele. Similarly, the dominance coefficient of the ancestral allele is $1 - h$ where $h$ is the dominance coefficient of the derived allele. The frequency of each allele is measured at the time of the shift in the male-female optimum. Alleles are divided into quartiles based on their average squared effect size across males and females ($\frac{1}{2}(e_m^2 + e_f^2)$). We observe the predicted negative relationship between dominance coefficient and frequency across the both X chromosome and the autosomes, particularly when alleles have larger average phenotypic effects and are therefore more 'visible' to selection.

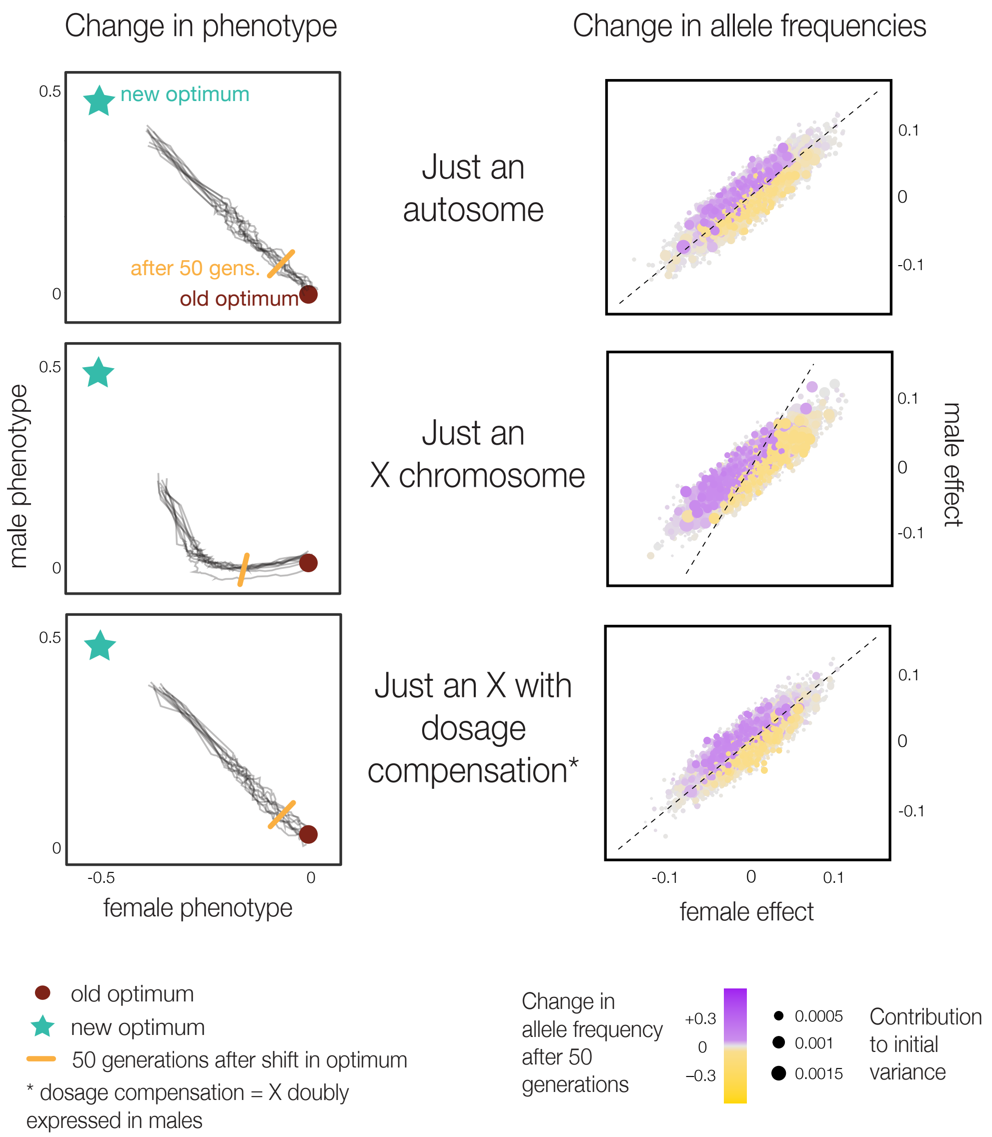


**Figure S5:** Same as Fig. 2, but male and female effect sizes are drawn from a bivariate normal distribution with variance $10^{-3}$ (instead of $10^{-2}$, as in the main text). Because allelic effect sizes are smaller, on average, relative to the magnitude of the shift in optimum, movement to the new male-female fitness optimum is slower across all genomic regions.

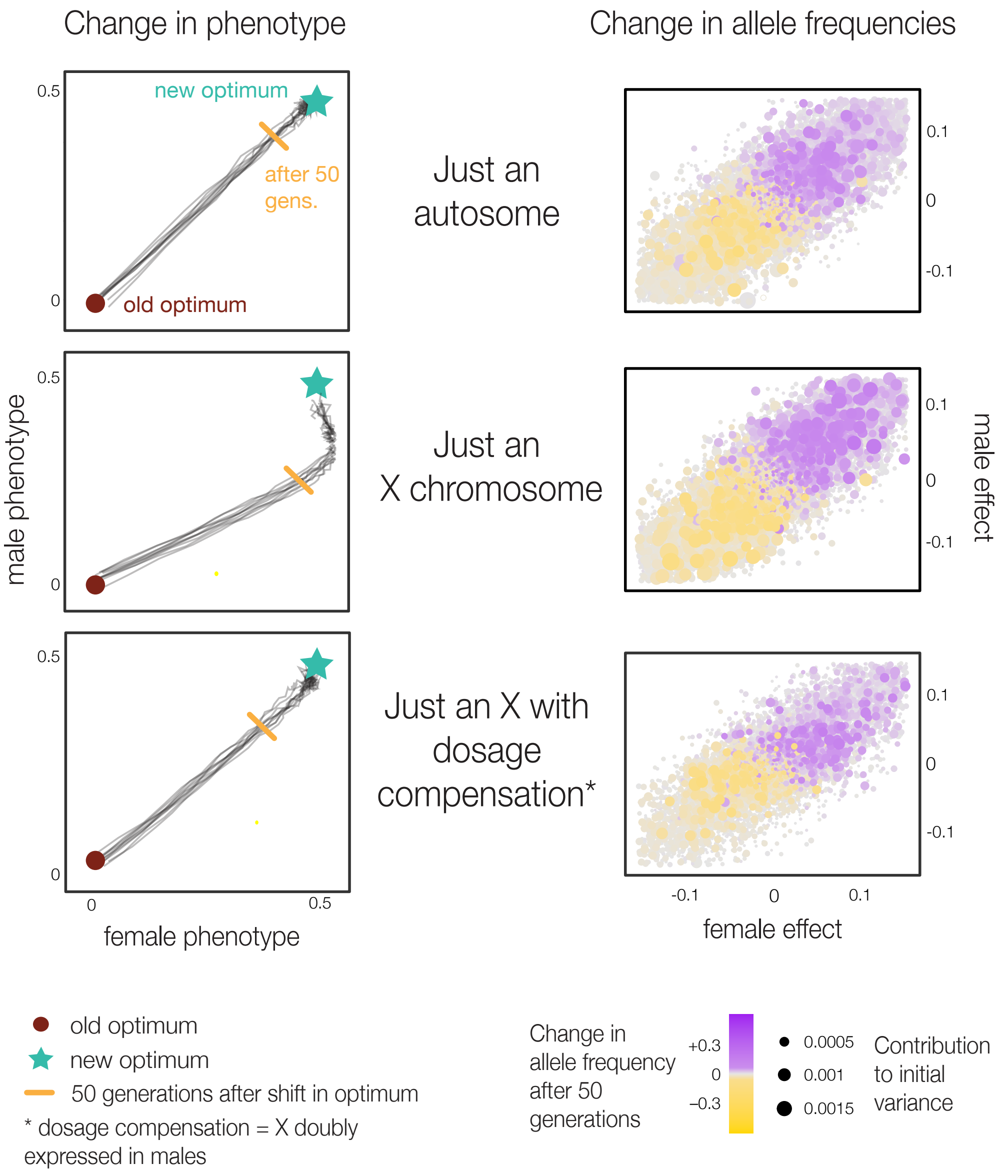


**Figure S6:** Same as Fig. 2, but the shift in the fitness optimum is in the direction of the male-female correlation of mutational effects. The magnitude of the shift in optimum is the same.

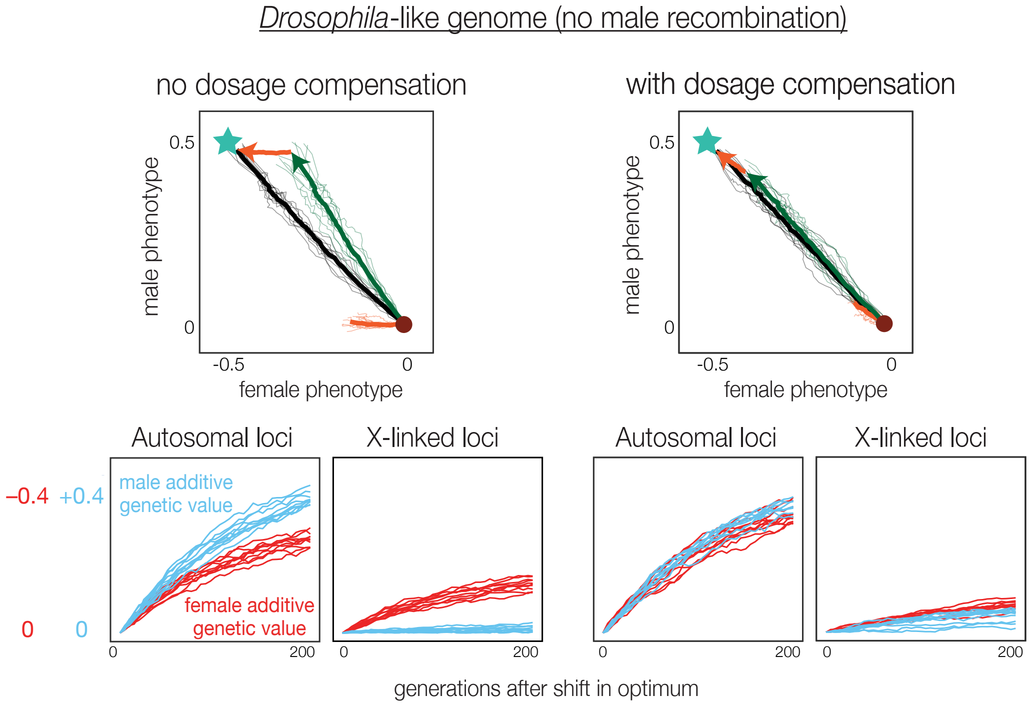


**Figure S7:** Same as Fig. 3, but with no autosomal recombination in males.

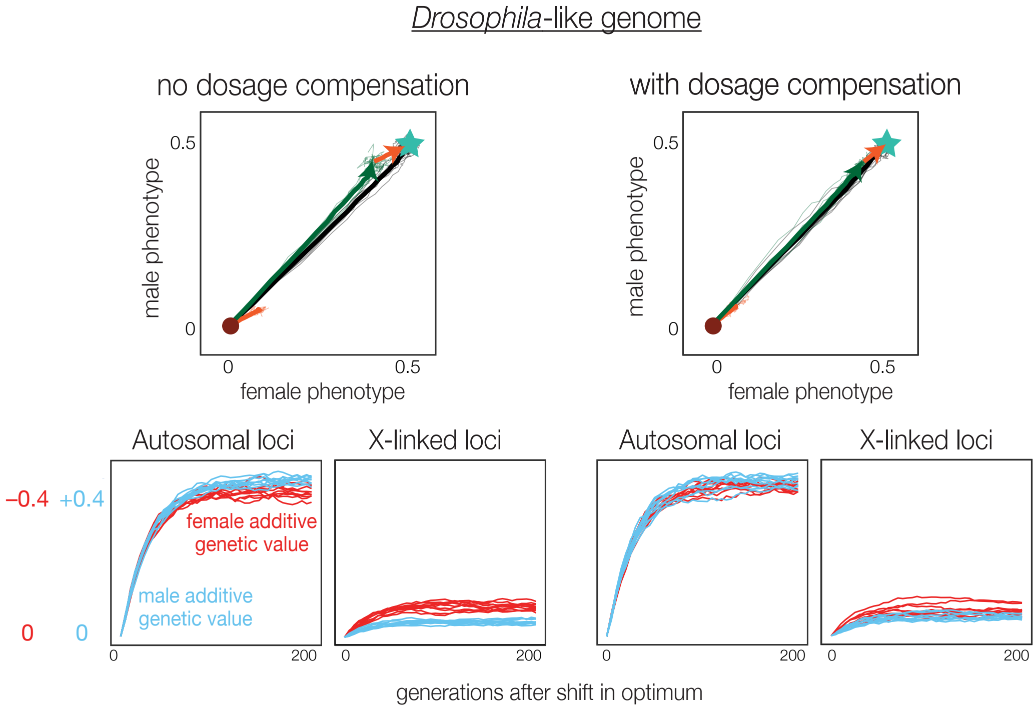


**Figure S8:** Same as Fig. 3, but with a concordant shift in the fitness optimum. The additive genetic value across X-linked loci still contributes more to movement to the female phenotypic optimum rather than the male, but this effect is reduced relative to the case of a discordant shift in the fitness optimum.

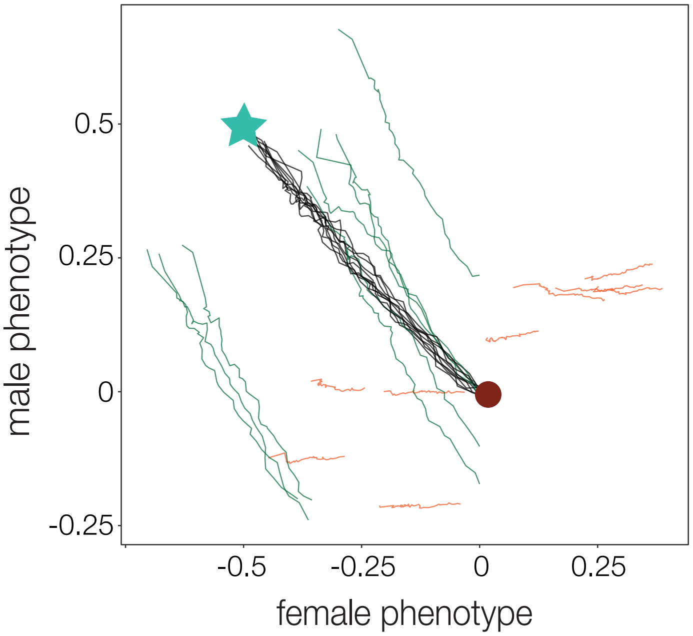


**Figure S9:** Same as Fig. 3, but without normalizing the starting additive genetic value to 0 for different genomic regions. While the overall starting phenotype across simulations is approximately 0 (black lines), this starting phenotype is achieved by different combinations of additive genetic values on the X chromosome and autosomes, which have diverged during the burn-in period of our simulations due to drift. While we do not track the additive genetic values on each autosome separately in these simulations, we would expect to see a similar pattern of diverged additive genetic values across different autosomes if we did so (i.e., the divergence of additive genetic values across the X chromosome and autosomes is not unique to these chromosomes)

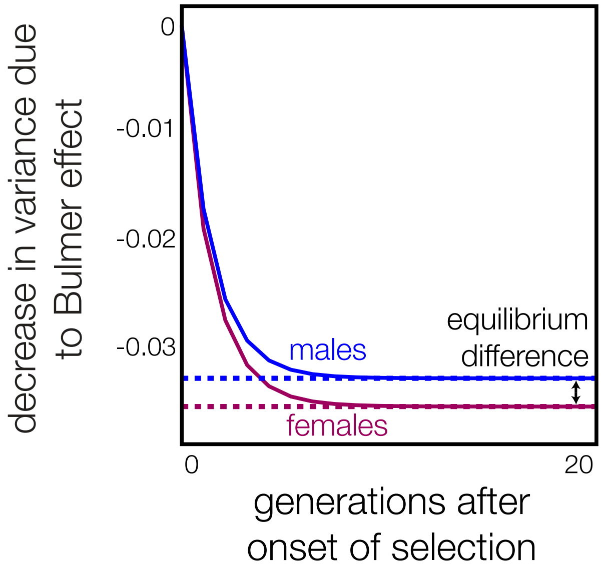


**Figure S10:** Same as Fig. 6B, but here males start out with a reduced phenotypic variance to reflect the haploidy of their X chromosome (Appendix).

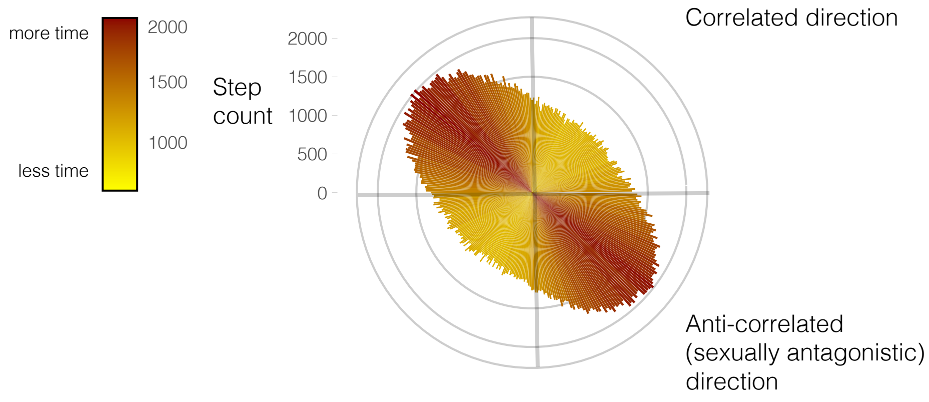


**Figure S11:** Same as in Fig. 7A, but here the male and female effect sizes of new mutations are drawn from a bivariate normal distribution with a correlation of 0.99 rather than 0.9. This stronger correlation of mutational effects drives a stronger genetic correlation between the male and female phenotype. A consequence of this stronger genetic correlation is that movement in a sexually antagonistic direction across an adaptive landscape is even slower than seen in Fig. 7A., because genetic variation is even more constrained to the correlated axis in male-female phenotype space.
